# Pattern of frustration formation in the functional brain network

**DOI:** 10.1101/2022.05.29.493932

**Authors:** Majid Saberi, Reza Khosrowabadi, Ali Khatibi, Bratislav Misic, Gholamreza Jafari

## Abstract

The brain is a frustrated system that contains conflictual link arrangements named frustration. The frustration as a source of disorder prevents the system from settling into low energy states and provides flexibility for brain network organization. In this research, we tried to identify the pattern of frustration formation in the brain at the levels of region, connection, canonical network, and hemisphere. We found that frustration formation has not a uniform pattern. Some subcortical elements have an active role in frustration formation, despite many low contributed cortical elements. Frustrating connections are mostly between-network types and triadic frustrations are mainly formed between three regions from three distinct canonical networks. Although there were no significant differences between brain hemispheres. We also did not find any robust differences between the frustration formation patterns of various lifespan stages. Our results may be interesting for those who study the organization of brain links and promising for those who want to manipulate brain networks.

## Introduction

The brain is an integrative system that its components cooperate to emerge advanced functions. So brain components should not always have seemed independent, but also sometimes must be regarded as a whole. In recent years, network modeling has been facilitating the study of the collective behavior of brain elements. Several classes of networks including simple connected-disconnected, weighted, and directed networks are used to clear brain mechanisms such as functional segregation and neural integration and help to discriminate brain disorders (Bassett & Sporns, 2017; Rubinov & Sporns, 2010; Mišić & Sporns, 2016; Bassett & Bullmore, 2009; Liu et al., 2017). Most of the works ignore the sign of links, either by taking the absolute value of connections or by positive thresholding on connection values (Garrison et al., 2015; Andellini et al., 2015; Wang et al., 2021; Theis et al., 2021). They minimize the impact of negative connections and disregard interactions between positive and negative connections.

Considering the brain as a signed network is also a promising approach to investigating the collective behavior of brain regions, where synchronous and anti-synchronous coactivations of brain regions determine positive and negative links, respectively (Saberi et al., 2021a) (Figure 1). Signed networks are mostly well-known for social system modeling in the context of friend and foe between entities (Tang et al., 2016; Yang et al., 2007; Yang et al., 2012; Facchetti et al., 2011). They are being investigated for the optimized arrangement of relationships using structural balance theory (Facchetti et al., 2011; Antal et al., 2005; Anchuri & Magdon-Ismail, 2012; Sheykhali et al., 2020; Derr et al., 2018). In this context, the optimized state happens when signed links have fewer higher-order conflictual relations and the network has less tendency to alteration and lower energy. In this way, we recently employed the theory to study the structural balance of the resting-state network (Saberi et al., 2021a). We found that minor negative brain connections get together in a way to make negative hubs to reduce the number of conflictual signed links arrangements which decreases brain energy and brings more stability to the brain network. Our results highlight the role of negative connections in brain network organization.

**Figure 1.**
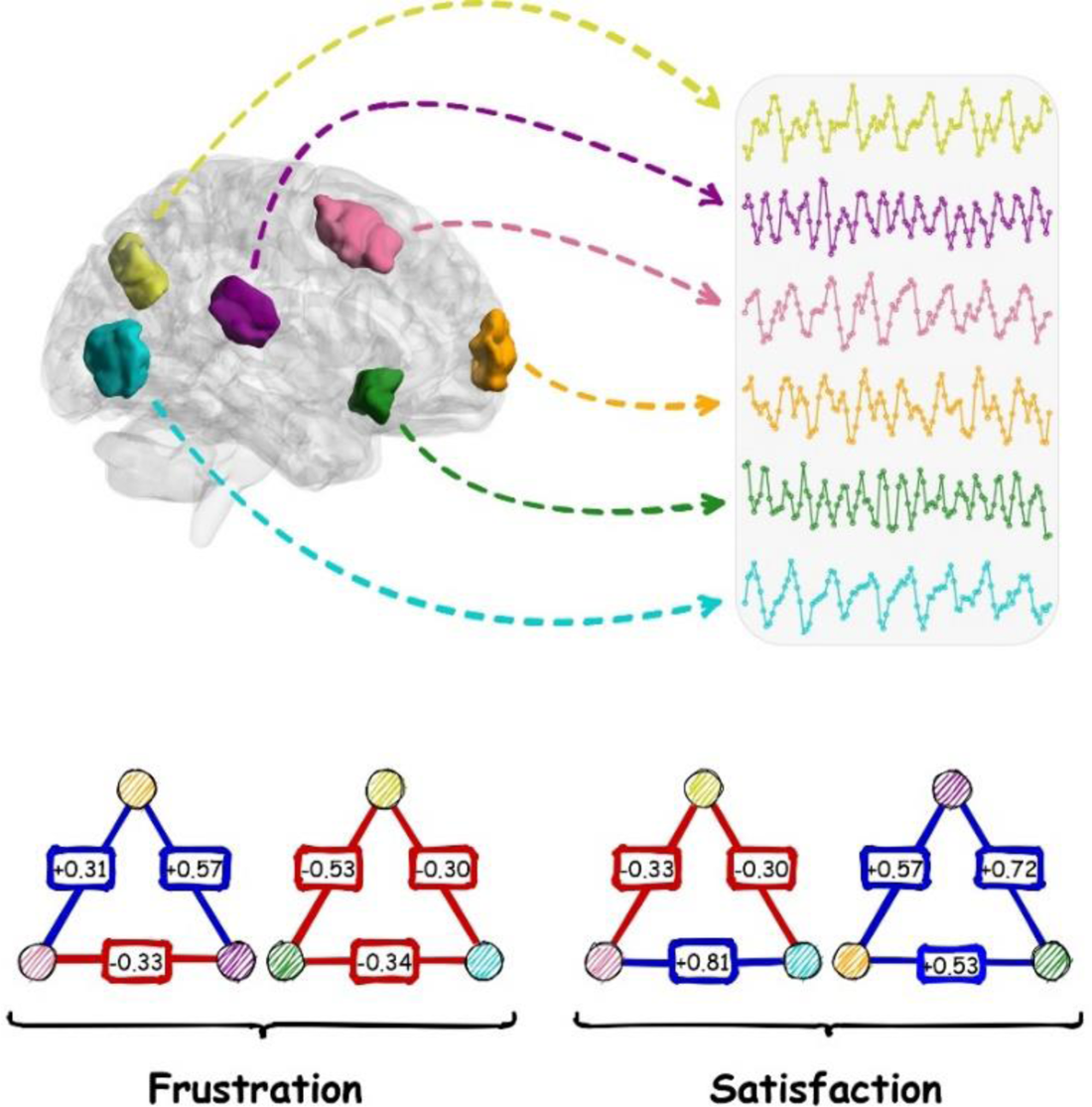
Formation of frustrated (imbalanced) and satisfied (balanced) triadic relations in the brain. Every region has a specific time course. Nodes of triads correspond to the regions by color. Correlation coefficients between every pair of activity patterns are also denoted on links. Blue and red links denote synchrony and ant-synchrony between time courses.

Frustration is another interesting phenomenon that can be studied in signed networks. The concept of frustration originates from the study of order-disorder systems in many-particle physics (Vannimenus & Toulouse, 1977; Toulouse, 1987, Villain et al., 1980) where it helps to understand the mechanism behind system ordering and phase transition (Zhao et al., 2019; Goremychkin et al., 2008). In spin systems, frustration is defined as topological constraints between spin neighbors that prevent minimizing system energy (Vannimenus & Toulouse, 1977). Generally, frustration as a source of disorder matters for systemic organization, alteration, and optimization.

In signed networks, frustration refers to non-trivial cycles of signed links, unstable assemblies that are seldomly found in real networks (Saberi et al., 2021a; Antal et al., 2005; Kirkley et al., 2019). Triadic relation is the smallest cycle where other elements influence the quality of a link, which makes a sense of the system (Winkler & Reichardt, 2013). It is analogous to the imbalance triads of Heider’s balance theory (Heider, 1946; Rapoport, 1963), “the friend of a friend is an enemy” and “the enemy of an enemy is an enemy”. The theory states that entities of nontrivial relationships are frustrated about their conditions and endure pressure to change the type of relationships to become balanced, “the friend of a friend is a friend” and “the enemy of an enemy is a friend”. These conflictual arrangements have been extended to any cycles with odd numbers of negative links (Cartwright & Harary, 1956; Estrada, 2019; Aref & Wilson, 2019).

Previous works showed that the resting-state brain signed network locates in a glassy state containing triadic frustrations (Saberi et al., 2021a; 2021b). These frustrations prevent the brain network to reach minimum energy (absolute stable state). In other words, the resting brain network is located nearly to transition to other states that highlights the role of brain network frustrations in the systemic reorganization of brain links.

Identifying brain network frustrations enables us to control the organization of brain links that affect the functionality and efficiency of the brain system. It’s helpful for those who are interested to induce functional changes by the use of neurostimulation and may promote understanding the mechanism of brain system functions and dysfunctions. So we decided to identify the pattern of frustration formation in the brain network system in the current study. In this regard, we investigated the contribution level of brain elements including regions, functional connections, canonical networks, and hemispheres in frustration formation. We explored them for each lifespan stages separately then compared the patterns of stages.

## Results

We designed this study to investigate the pattern of frustration formation in the brain. So we explored how brain components contribute to brain network frustrations. We preprocessed anatomical and resting-state fMRI images of healthy subjects from two online repositories of ABIDE (Di Martino et al., 2014) and Southwest University (Wei et al., 2018) which can ensure the reliability of the results. Then we extracted regional activations of each functional image based on the parcels of Shen’s atlas (Shen et al., 2013) where the regions were projected to canonical brain networks (Yeo et al., 2011) (Figure 2). After that, we calculated functional connectivities and formed a signed network for each subject based on the signs of connections. Then we identified the triadic frustrations of each network and measured the contribution level of brain regions, functional connections, canonical networks, and hemispheres in their formations. On another side, we estimated the null contribution values of mentioned elements according to the number of appeared frustrations for each subject’s network. Finally, we performed a group-level paired comparison for the contribution of every mentioned elements between actual and null values to find out which regions, connections, canonical networks, and hemispheres were significantly involved in the frustration formation for each stage separately (Figure 3-6). In the end, we would have provided a multimodal map for frustration formation in the brain stagewise and without considering stages. We also investigated any significant differences between contribution patterns of lifespan stages.

**Figure 2.**
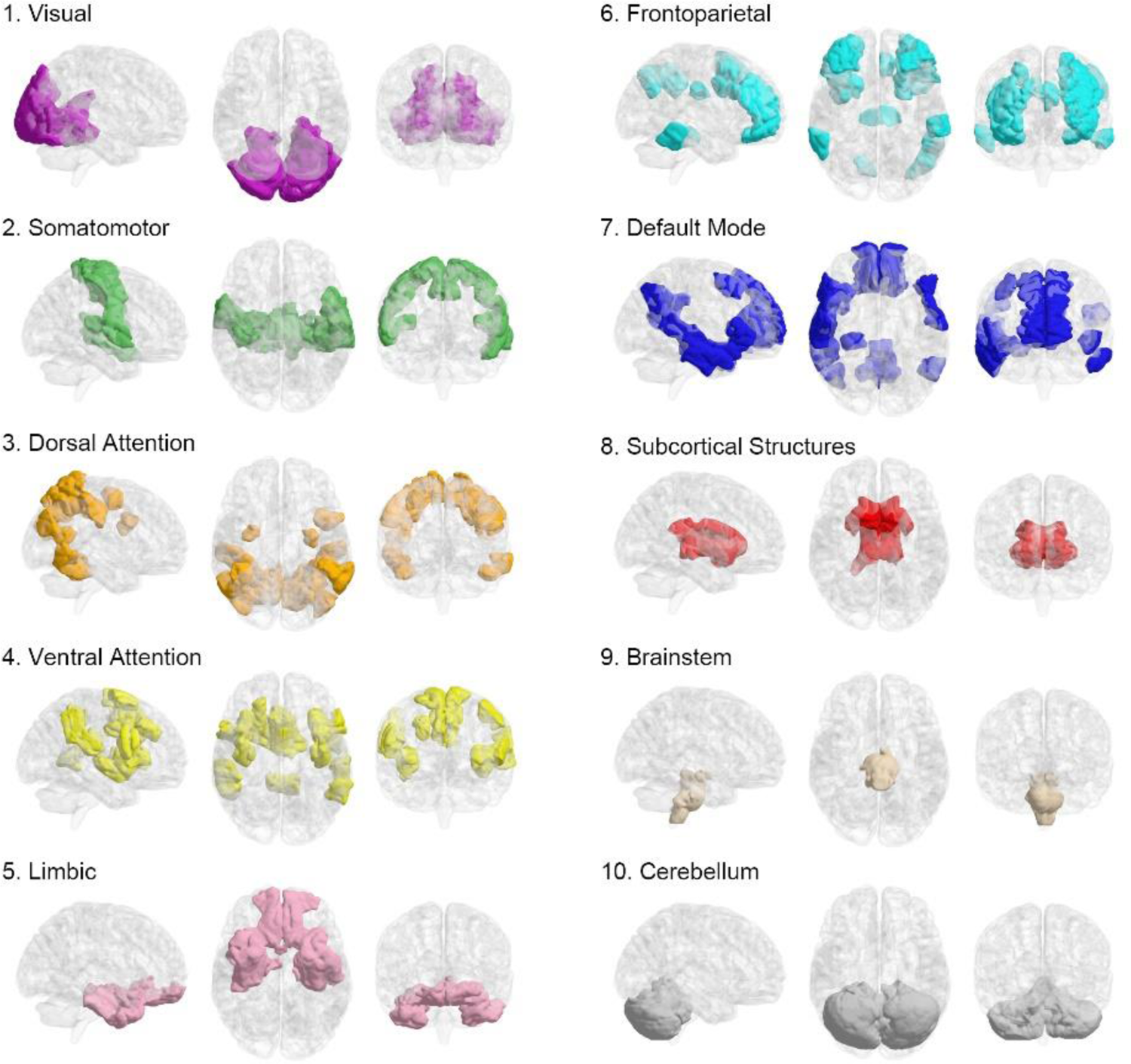
Categorizing Shens’ regions of interest into canonical networks.

**Figure 3.**
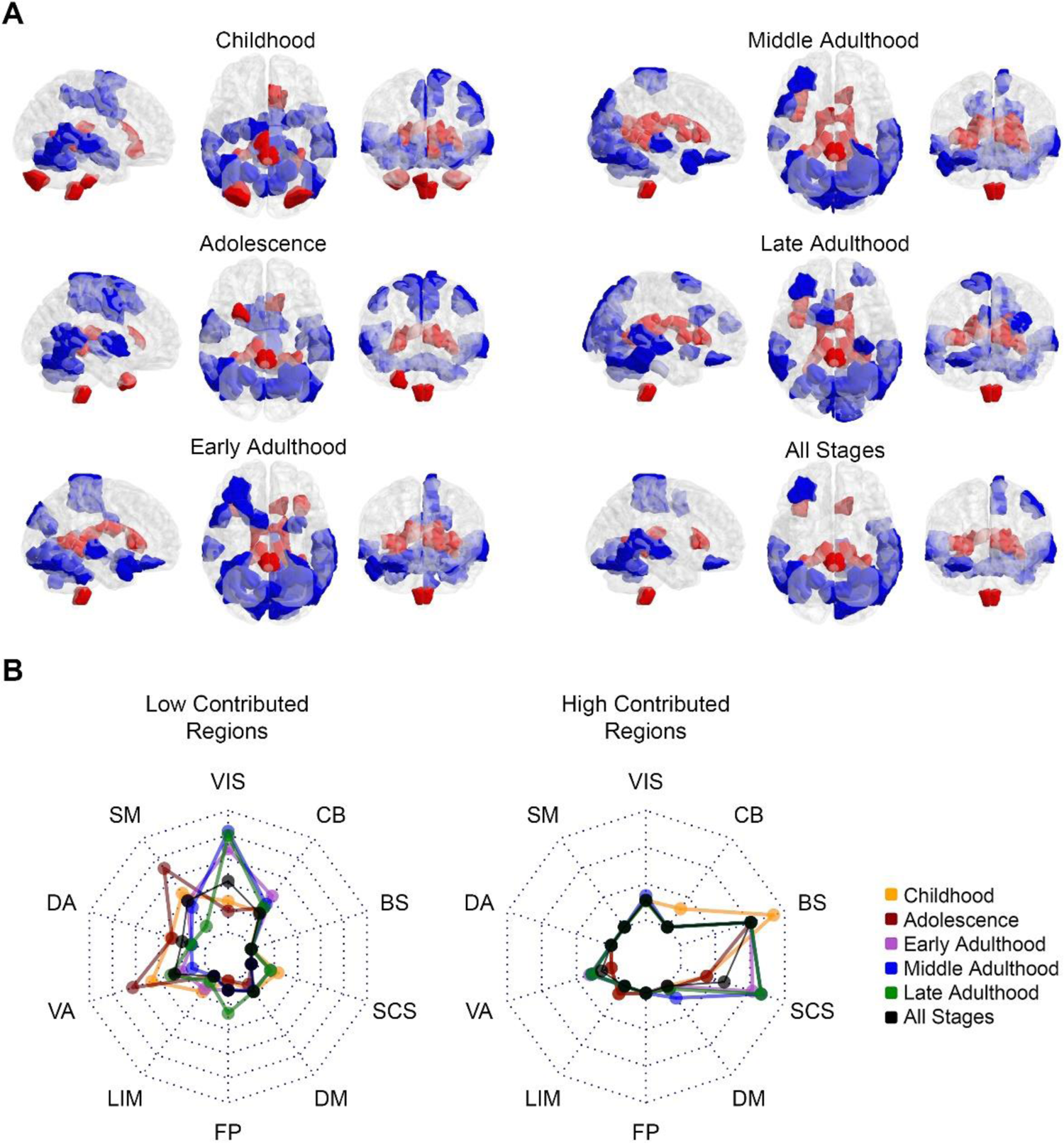
Pattern of regional contribution in frustration formation. **(A)** Red-colored and blue-colored areas of the brain maps indicate Shen’s regions with significantly greater and significantly lower contributions in frustration formation. Brain maps demonstrate the patterns for various lifespan stages and all stages in three representational planes of sagittal, axial, and coronal. **(B)** Radar charts show what percentage of canonical brain networks are involved in significant areas of (A). The left radar plot relates to blue areas and the right one corresponds to red areas. Each dashed polygon indicates 10 percent. Various line colors are related to lifespan stages. Abbreviation; VIS: visual, SM: somatomotor, DA: dorsal attention, VA: ventral attention, LIM: limbic, FP: frontoparietal, DM: default mode, SCS: subcortical structures, BS: brainstem, CB: cerebellum.

### Contribution of brain regions

For each subject, we recognized presented frustrations and determined the number of frustrations that each region is involved in them. On the other hand, we calculated the expected values of regional involvements in the case of a uniform engagement of regions in frustration formation. The expected contribution value of a region of interest (ROI) for a subject’s whole-brain network with N_ROI_ regions and N_Frust_ frustrations is derived as follows:

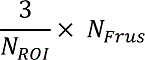

Number 3 is multiplied since a frustration occurs between three regions. In this way, we provided actual and null values of contributions in frustration formation for 268 regions of each subject. Then we performed a paired group-level comparison between actual and null values of each region to find which regions have a significantly different contribution. We analyzed subjects of each stage separately in addition to disregarding the lifespan stages (Figure 3). Since contribution values were distributed non-normally, we used Wilcoxon matched-pairs signed-rank test for this goal. Figure 3A indicates regions with significantly lower (blue-colored) and significantly greater (red-colored) involvements in frustration formation compared to expected values after multiple comparison corrections on p-values using False Discovery Rate (FDR) and with large effect sizes (greater than 0.6). We reported the corrected p-values and effect sizes in supporting information. Since the frustration expresses disorganization in the network, the blue and red areas of the figure represent regions that have well-organized and disorganized relations with other regions, respectively. To better interpret the results, we mapped significant brain regions to canonical networks of Figure 2 to show how red and blue areas belong to the canonical networks. Method section “parcellation atlas projection” describes the projection process that we used to obtain Figure 2 in detail. Radar plots of Figure 3B represent what percentages of canonical networks have significant contributions where dotted polygons divide the whole network volume by ten percent. The left and right radar charts correspond to blue-colored and red-colored regions of Figure 3A. The left radar chart shows low contributed blue regions mostly belong to the visual network in adulthood and they involve somatomotor and ventral attention networks in the early stages. More than 40% of the visual network in middle adulthood and about 35% of the somatomotor network and ventral attention network in adolescence have regions with well-organized relations to other brain regions. The right radar chart also shows that most of the high contributed red regions of Figure 3A belong to subcortical structures and the brainstem. Despite early stages, more than 20% of subcortical structures in adulthood have areas with disorganized relations to other regions. Also, more than 20% of the brainstem volume is significantly involved in frustration formations, it seems an inferior portion of the brainstem according to Figure 3A. Radar charts also indicate moderate values for the all stages brain map, which we desired in case of disregarding stages. We also used the nonparametric Kruskal-Wallis test to investigate between stage differences of regional contributions. We performed the test between contribution values of various stages for each region separately and then corrected p-values from false positives that occurred by multiple comparisons using FDR besides effect size measurement. There is no region with a significant corrected p-value and large effect size (greater than 0.14) that indicates no robust contribution differences between stages. Although Figure S1 shows regions with corrected p-value lower than 0.05 and medium effect size (between 0.06 and 0.14). To test whether significant regions of Figure 3A appear randomly or not, we compared the entropy of the all-stage pattern with the entropy of its shuffled pattern for greater and lower contributions separately. I explained the way we calculated entropies in Method section of “entropy calculation”. Figure S2 shows the histogram of entropies related to shuffled patterns compared to the entropy of actual patterns. It indicates that both lower contributed regions (blue-colored) and greater contributed regions (red-colored) have significantly lower entropies which validates the non-randomness of regional significant patterns of Figure 3.

### Contribution of functional connections

After regional investigations, we explored the contribution of functional connections in frustration formation. So for each subject, we counted the number of triadic frustrations that each connection is involved in them. Also, we estimated the expected contribution values of each connection in case of uniform involvement of connections as follows:

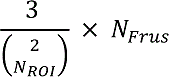

Where *N_Frust_* and *N_ROI_* are the number of presented frustration and the number of nodes in the subject’s network. 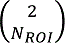 also denotes the 2-combinations of *N_ROI_* that is equals to the number of links in a fully-connected network with *N_ROI_* regions. The factor of 3 is also appeared because a frustration engages three connections. In this way, we calculated actual and null contribution values for each connection of each subject. After that, we performed a paired group-level analysis between actual and null values for each functional connection using Wilcoxon matched-pairs signed-rank test. We chose this non-parametric test since the values were not distributed normally. Figure 4 displays contribution maps of functional connections in frustration formation for each stage separately and all stages. Each cell corresponds to a functional connection between two regions of interest, so heatmaps represent mutual connections between 268 Shen’s regions. The regions are categorized based on networks of Figure 2 by segmenting the axes of heatmaps. Also, black squares discriminate between-network connections. Blue and red cells indicate connections with significantly lower and significantly greater contributions in frustration formation that have FDR corrected p-values smaller than 0.05 and large effect sizes (greater than 0.6). The figure does not exhibit a distributed pattern of high contributed connections (red-colored cells). It seems that connections of several visual and subcortical regions with other regions have significantly great contributions to frustration formation. Many default mode to ventral attention connections of adulthood are also high contributed. On the contrary, lower contributed connections are more distributed in the brain network. Most of these connections are within-network types, especially those located in visual, somatomotor, and attentional networks. There are also lots of between-network connections that have lower contribution in frustration formation, especially connections between visual, somatomotor, cerebellar, and attentional regions. We reported corrected p-values and effect sizes of the comparisons in supporting information.

**Figure 4.**
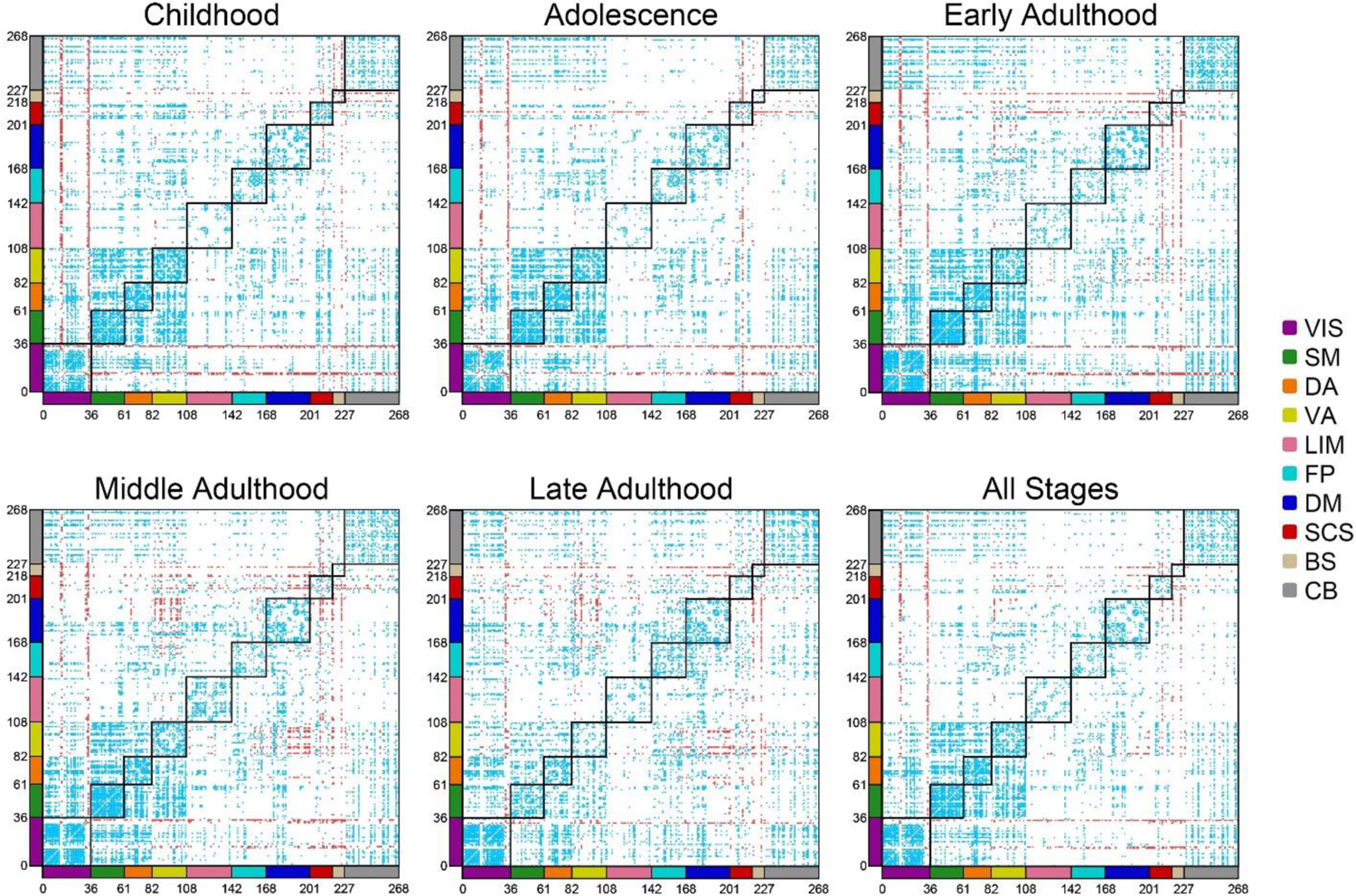
Contribution maps of functional connections in frustration formation. The first five maps demonstrate the maps related to lifespan stages and the last one corresponds without considering the stage. Shen’s 268 ROIs are categorized into 10 networks of Figure 2 by axes coloring. Cells also represent functional connections between every two ROIs. Red cells display those connections with a significantly greater contribution to frustration formation and blue cells indicate those connections with significantly lower involvement. Black squares discriminate between network connections. Abbreviation; VIS: visual, SM: somatomotor, DA: dorsal attention, VA: ventral attention, LIM: limbic, FP: frontoparietal, DM: default mode, SCS: subcortical structures, BS: brainstem, CB: cerebellum.

As well as the last section, we also wanted to explore the randomness of the pattern of the significantly contributed connections. Based on what we described in the Method section, we estimated the entropy of the all-stage subfigure of Figure 4 and its shuffled patterns for significantly lower connections (blue-colored) and significantly greater connections (red-colored) separately. Figure S3 shows that the pattern of blue-colored connections and the pattern of red-colored connections have lower entropies compared to shuffled patterns that state significant connections are not random.

In addition, a question has arisen whether high contributed and low contributed connections that are denoted in Figure 4 are within-network types or between-network types. To answer this question and better interpret the heatmaps of Figure 4, we created a metric and named it WBR. WBR measures the ratio of within-network connectivity to between-network connectivity. As such that we first consider our interesting connections, for example higher contributed connections, then count the number of appearing between-network connections and within-network connections. On another side, we calculate the number of possible between-network connections and within-network connections based on the number of nodes. Finally, we divide the calculated values as below:

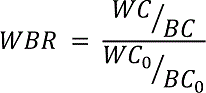

Where *WC* and *BC* stand for the number of within-network connections and the number of between-network connections and *WC_0_* and *BC_0_* are also their maximum possible values. *WBR* would be more than one in the case of intense within-network connections, lower than one in the case of extreme between-network connections, and equal to one in the balance between them. Table 1 demonstrate *WBR* of high contributed connections and low contributed connections in frustration formation calculated based on heatmaps of Figure 4. The results show that the low contributed connections tend to be within-network types and high contributed connections are mostly between-network types. We also investigated the difference in the contribution of connections in frustration formation between lifespan stages. So we performed a multi-group Kruskal-Wallis test for each connection between observed contribution values of different stages subjects. Then we corrected multiple comparison effects on p-values using the FDR method. We did not find any significant corrected p-values with large effect sizes (greater than 0.14) for various functional connections. Figure S4 only shows connections with corrected p-values lower than 0.05 with medium effect sizes (between 0.06 and 0.14)

**Table 1.**
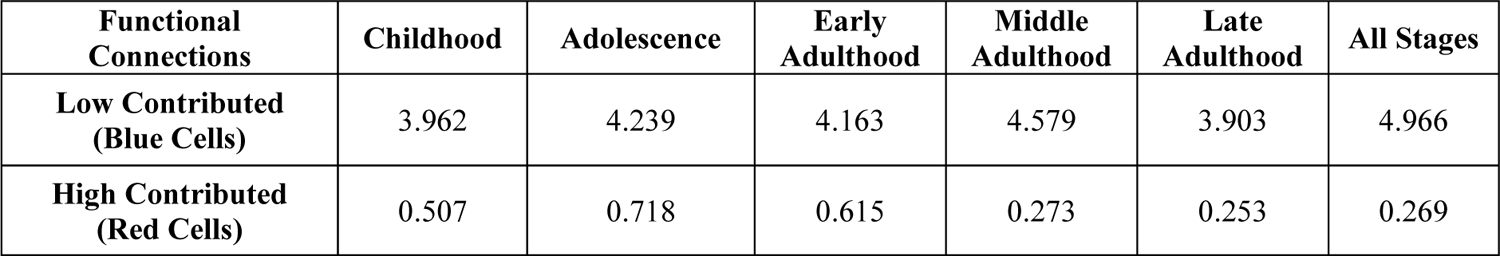
WBRs calculated based on low contributed and high contributed connections of **Figure 4** heatmaps.

### Contribution of canonical networks

In the last sections, we explored the involvement of brain regions and their functional connections in frustration formation. In this section, we wanted to study the contribution of canonical networks. So we used the projection of Shen’s ROIs into Yeo’s networks (Figure 2 and method section “parcellation atlas projection”). There are 4 types of frustration formations that regions of a canonical network can be involved in them (Figure 5), 1) within-network: all three regions belong to the target network, 2) between-network (I): two regions located in the target network and one another outside, 3) between-network (II): one region located in the target network and two other regions in another network, 4) between-network (III): one region located in the target network and two other regions in two other distinct networks. For every combination of formation type, stage, and canonical network, we performed a pairwise group comparison using the Wilcoxon signed-rank test between actual and expected contribution values of subjects and then integrated all results into Figure 5 and Tables S1-S4. Formulas to calculate null contribution values of the target network of *T* in mentioned frustration formation types are described as follows:

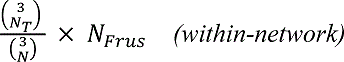

Where *N* and *N_T_* denote total ROIs number and those ROIs number located in canonical network of 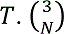 and 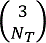 equal to the possible triangle numbers in whole-brain network and the possible triangle numbers in the target network. *N_Frus_* is also the number of appeared frustrations in corresponding actual whole-brain network that we wanted to calculate its null values.

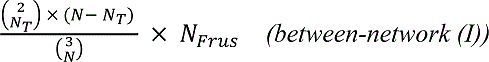

The numerator of the fraction is equaled to the number of triangles that their two ROIs located in the target network of *T* and one another outside.

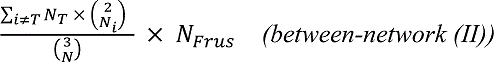

**Figure 5.**
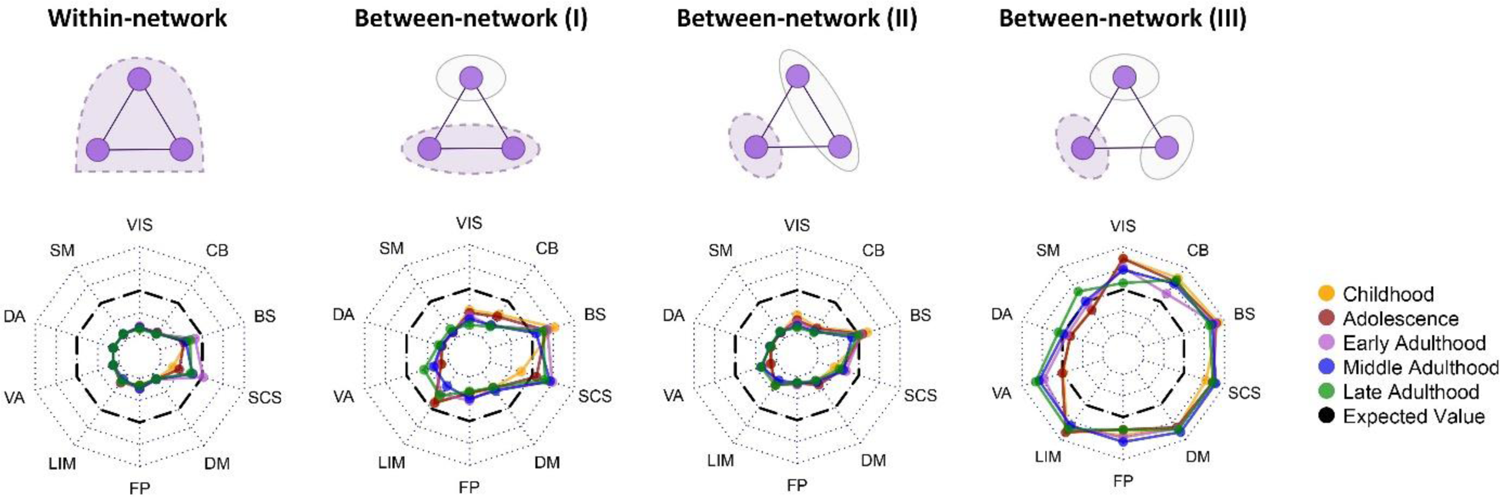
Contribution of canonical networks in frustration formation. The above row displays 4 possible frustration formation types between regions of canonical networks, from left to right: all regions located in the network, two regions in the network and another outside, one region in the network and two other regions in another network, one region in the network and other regions in two distinct networks. Dashed violet binds denote the elements of interest in analysis. Radar charts correspond to every frustration formation type displayed above them and show the difference between actual and expected contribution values. Dotted lines of radar charts segment r effect size from −1 to 1 where black dashed lines indicate zero r effect size. Line colors demonstrate different lifespan stages. Effect size; small: r < 0.4, medium: 0.4 < r < 0.6, large: r > 0.6. Abbreviation; VIS: visual, SM: somatomotor, DA: dorsal attention, VA: ventral attention, LIM: limbic, FP: frontoparietal, DM: default mode, SCS: subcortical structures, BS: brainstem, CB: cerebellum.

The summation performs over all the networks except network of *T* and the numerator of the fraction is equaled to the number of triangles that their one ROI located in the target network and two ROIs in another super-region.

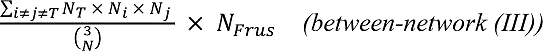

The summation performs over every distinct pair of networks except network of *T*. The numerator of fraction denotes possible triangles that their one ROI located in the target network and two other ROIs belong to two other distinct networks. After counting contribution values for formation types and calculating null values, we performed paired group analysis. All the effect sizes and corrected p-values are reported in supporting information. Radar charts of Figure 5 demonstrate the r effect size of Wilcoxon matched-pairs signed-rank tests between actual and null contribution values. The results demonstrate that all networks in all of the stages except all stages’ brainstem and adults’ subcortical structures have significantly lower within-network types of frustration formation with large effect sizes (greater than 0.6). Visual, somatomotor, dorsal, and ventral attention, frontoparietal, and default mode networks have significantly lower contributions in between-network frustration formation type I, although their effect sizes are medium (between 0.4 and 0.6) in some stages. Also, early adults’ limbic networks and adults’ cerebellums have significantly lower contributions in between-network type I and large effect sizes. Subcortical structures in early and middle adulthood and brainstem in childhood have significantly higher contributions in the formation of between-network type I with medium effect sizes. Investigation of between-network type II also shows that most of the networks in all stages have significantly lower contributions in frustration formation with large effect sizes except brainstem and subcortical structures where subcortical structures just in childhood, adolescence, and late adulthood have medium effect sizes. The last radar chart (between-network III) shows that most of the higher contributions occur between 3 distinct networks. Although it indicates that somatomotor and dorsal attention are not contributed to this type of frustration formation. Almost, other networks of all stages have significantly higher contributions in this type of frustration formation with large or medium effect sizes except visual network in late adulthood, ventral attention in childhood and adolescence, and frontoparietal in late adulthood that have not significant p-values, and frontoparietal in adolescence and cerebellum in early adulthood that have small effect sizes. We also compared contribution values of different stages for every pair of formation types and canonical networks using the Kruskal-Wallis test. The test obtained no significant corrected p-values with large effect sizes. P-values and effect sizes of these analysis are also presented in supporting information.

### Contribution of hemispheres

In the end, we investigated the contribution of the brain hemispheres in frustration formation. There are 4 possible states for the contribution (Figure 6A): 1) all three regions of frustration belong to the right hemisphere, 2) all three regions belong to the left hemisphere, 3) only two regions located in the right hemisphere, 4) and only two regions located in the left hemisphere. We compared actual and null contribution values for every combination of state and stage using the Wilcoxon matched-pairs signed-rank test. We estimated the expected contribution values of 4 states for a network as follows:

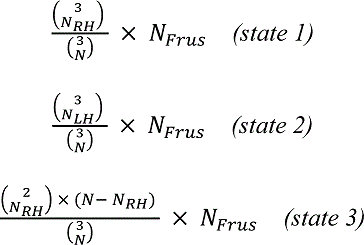

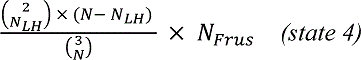

where *N_RH_* and *N_LH_* are the number of regions in right and left hemispheres that are equaled to 133 and 135 according to the Shen’s atlas. Figure 6B shows the result of comparisons. We reported their statistics in Table S5. The results show no significant corrected p-values with large effect size (greater than 0.6) for the comparisons. All effect sizes were small except state 4 in middle adulthood that has medium effect sizes (r effect size = 0.408). We also compared contribution values of different stages in every state and did not find any non-small effect sizes for between-stage differences. To investigate the main effect of the hemisphere on frustration formation disregarding lifespan stage, we also performed two Wilcoxon matched-pairs signed-rank tests, one between state 1 and state 2 contribution values and the other between state 3 and state 4 contribution values (Figure 6C). Only the comparison between state 1 and state 2 had a significant p-value although its effect size was small (r effect size = −0.38). Generally, we did not find any robust hemispherical effect on frustration formation.

**Figure 6.**
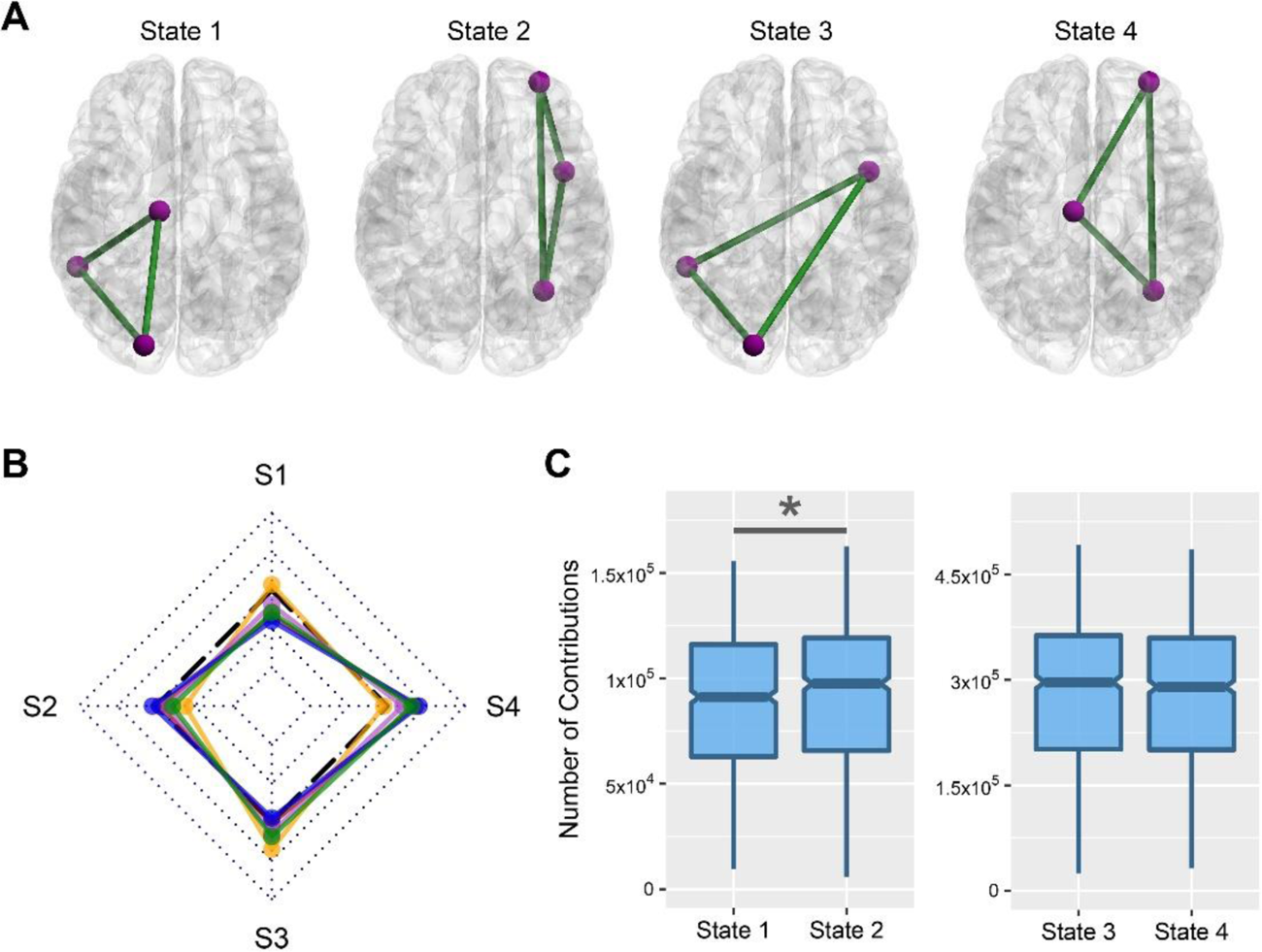
Contribution of brain hemispheres in frustration formation. **(A)** Four possible frustration formation types between brain hemispheres. **(B)** The difference between actual and null contribution values of every stage and state. Dotted rhombuses segment r effect sizes from −1 to 1 where black dashed lines denote zero value. S1-S4 corresponds to the state1-state4. Lines colors; orange: childhood, red: adolescence, violet: early adulthood, blue: middle adulthood, green: late adulthood. Effect size; small: r < 0.4, medium: 0.4 < r < 0.6, large: r > 0.6. **(C)** Comparison between the contribution of hemispheres without considering the stage. Horizontal lines of boxes indicate medians and notches determine the 95% confidence interval for the medians. Asterisk denotes the significant corrected p-value with a small effect size.

## Discussion

We wanted to find out how brain elements involve in frustration formation. So we compared the number of frustrations that each element is contributed to them with the null contribution value of that element. The null values are estimated based on the hypothesis of the uniform contribution of the elements in frustration formation. We performed the comparison in four levels of the brain region, functional connection, canonical network, and hemisphere. We did it for each lifespan stage separately as well without considering stages. We also compared the involvement of different stages. Regional level comparisons show that some brain regions have greater contributions and some regions have lower contributions in frustration formation, although we did not find any robust differences between the regional contributions of different stages (Figure 3). Investigation of functional connections also indicated that many functional connections have significantly lower or greater contributions to frustration formation (Figure 4). Low contributed connections are mostly within-network type and high contributed connections are usually the between-network type (Table 1). We also did not find any robust variation in the contribution of connections between stages. In addition, the results show the most significant regions and connections are lower contributed. Entropy analysis also indicated that regional and connectional significant patterns are not random. In the following, we studied the contribution of canonical networks in frustration formations. We found that most frustrations appear between 3 regions of 3 distinct networks (Figure 5) and there are no powerful differences between network-based contributions of lifespan stages. We also did not find any strong hemisphere related effects on frustration formation, no difference between the role of the right and left hemispheres (Figure 6).

### Role of subcortical structures

We obtained that subcortical regions have a prominent role in frustration formation in the brain network at both nodal and connectional levels. So It can emerge instability and altering properties that facilitate systemic level neural changes and bring adaptive characteristics to the brain. Certainly, we can not understand brain reorganization without taking into account the white matter tract located in subcortical areas. Older prospectives considered subcortical alterations as a passive consequence of cortical functional reshaping however new mechanisms underline the active reorganization of sub-circuits within the large network of the brain (Duffau, 2009). Animal studies posited brainstem-related and subcortical plasticity under exposure to visual and auditory stimuli (Duménieu et al., 2021; Chandrasekaran et al., 2014; Miranda et al., 2014). Human studies reported such subcortical rewiring mechanisms for skill learning such as second language and motor training (Lui et al., 2020; Sampaio-Baptista et al., 2013; Scholz et al., 2009). A recent study also indicates subcortical short-term plasticity yielded by deep brain stimulation pulse (Awad et al., 2021). All these observations are agreeing with the frustrating essence of the brain subcortex. In addition, we know that many studies reveal malfunction and abnormal structural changes in the subcortex of neural disorders (Arnold Anteraper et al., 2014; Cerliani et al., 2015; Lee et al., 2018; Rosenberg-Katz et al., 2016; Hoogman et al., 2017) while subcortical small changes may have extensive effects at the cortical level (Jones et al., 2000). So controlling the activation of frustrating subcortical regions can provide the favorable systemic reorganization of the brain network. It may be appliable by focusing on our detected frustrating regions and connections using sophisticated subcortical stimulation methods (Awad et al., 2021; Colle et al., 2021; Folloni et al., 2019; Shi et al., 2021). In addition, the prominent role of the subcortex in frustration formation may be related to its unidirectional projections (Shi et al., 2014) or indirect pathways (Lanciego et al., 2012) which need more exploration.

### Between-network frustrations

The brain is a self-organized system (Dresp-Langley, 2020) such that functional signed links adopt a topology to reduce network frustration (Saberi et al., 2021a). We found that this property is consistent within canonical networks as well where canonical networks have lower numbers of frustrations compared to null (Figure 5). Most of the frustrating connections are formed between networks (Table 1) and frustrated triangles mostly engage three distinct networks (Figure 5). This is what we expect to happen since large-scale brain networks consist of coactivating local brain seeds (Yeo et al., 2011). They decrease the chance of negative functional connection appearance and subsequently frustration formation. So if someone is interested in handling brain network frustration he/she should pursue a multi-network approach.

### Visual network contribution

Our results indicate that most visual regions of adults have a low contribution to frustration formation (Figure 3) which may be a consequence of visual maturation (Siu & Murphy, 2018). Although several ROIs belonging to the hippocampus show frustrating connections to most brain areas (Figure 4). It’s so fascinating result since the prominent role of the hippocampus in learning and memory and its association with broad areas of the brain to play this role (Anand & Dhikav, 2012) is consistent with the frustrating essence of its connections. We mention that frustration facilitates adaptive processes due to their unstable nature.

### Somatomotor network contribution

As Figure 3 shows nearly 40% of somatomotor and ventral attention regions of adolescents have a lower contribution to frustration formation than adults. We wonder why this happens while neural flexibility is an essential developmental feature of adolescence. So we could not find a proper answer and any biological relevance for this effect.

### Ventral attention to default mode connections

We also saw some ventral attention to default mode connections are frustrating (Figure 4). This effect is enhanced in the adulthood range. We could not find a good psych-neural interpretation for this observation. Although a recent paper highlighted connectivity between ventral attention and default mode areas in Bulimia Nervosa (Domakonda et al., 2019). Because of the high comorbidity of Bulimia Nervosa by other mental disorders such as depression and bipolar (Walsh et al., 1985; McElroy et al., 2005) and the frequency of mood disorders in adulthood range, we think the frustrating properties of these connections may be related to these types of disorders.

### Lifespan comparisons

In one of our previous works when we compared the number of frustration between lifespan stages to track the requirement to change of brain network we reported a significant difference between stages (Saberi et al., 2021b). Because of the large sample size of stages, we decided to consider effect size besides corrected p-value in current work comparisons to specify more robust differences. Therefore we did not find any strong differences between frustration formation patterns of various stages, in other words, all significant comparisons with corrected p-value lower than 0.05 had corresponding small and medium effect sizes.

### Shen’s atlas to Yeo’s atlas projection

As we wanted to investigate the quality of canonical brain networks’ contribution in frustration formation we needed projection of Shen’s 268 ROIs into them. Finn et al. (2015) utilized Shen’s clustering algorithm to group the ROIs into eight networks of the medial frontal network, frontoparietal network, default mode network, subcortical and cerebellar regions, motor network, visual I network, visual II network, and the visual association network (Shen et al., 2013; Finn et al., 2015). The labeling is presented on “https://bioimagesuiteweb.github.io/webapp/connviewer.html<u>”</u> and has been used for the between-network analysis of studies (Rosenberg et al., 2016; Chen et al., 2020). When we explore the output of the categorization we can observe much implausible labeling for instance; some superior regions labeled as subcortical and cerebellar networks, some temporal regions categorized into medial frontal, and some cerebellar regions classified as frontoparietal, visual, and default-mode networks. Also, the limbic network which is one of the most important functional brain subsystems is not regarded. Since three networks are nominated based on the sense of vision, a question has arisen if the sensory system has been considered a major element of the labeling and why there is no sign of other sensations such as auditory in the ROIs categorization. As it seems some shortages in the categorization process, we decided to use a new standard procedure (Meyers et al., 2019) to project Shen’s ROIs into Yeo’s 7 large-scale brain networks (Yeo et al., 2011) based on the coincidence index of the two atlases (Figure S5). Since Yeo et al. (2011) excluded subcortical areas we specified them into three distinct networks according to their anatomical characteristics. We described the procedure and the intermediate results in the method section of “parcellation atlas projection”. So our optimized model classified Shen’s regions into 10 subnetworks of visual, somatomotor, dorsal attention, ventral attention, limbic, frontoparietal, default mode, subcortical structures, brainstem, and cerebellum (Figure 2).

### Global signal effects

There is much conflictual evidence on the quality of removing global brain signals, some studies suggest this step in pre-processing and some others decline it (Aquino et al., 2020; Murphy et al., 2009; Fox et al., 2009; Liu et al., 2017; Schölvinck et al., 2010). It’s clear that global signal regression produces such anti-synchronous connections with negative correlations (Figure S6) that matter for signed networks and can affect the structural balance (Fox et al., 2009). It’s also a fact that increasing negative links grows the number of frustrations since they have more negative links as compared to satisfaction (Figure 1). In the analysis of the main manuscript as well as our previous research (Saberi et al., 2021a; 2021b), we ignored global signal regression and explored the formation of frustration in the presence of minor negative links. Besides, we performed the same analysis on the global signal regressed functional images to check the difference. Figure S7-S8 shows the result of the analysis for the contribution of regions and contribution of connections in frustration formation. Figure S7 shows a lower number of significant regions compared to Figure 3 and indicates some other regions. Low contributed regions belong to somatomotor and attentional networks in adulthood and default mode in all stages and high contributed regions are mostly located in the subcortex in early adulthood and outspread in other stages. Investigation of connection contribution on the regressed images (Figure S8) also shows fewer numbers of significant connections compared to Figure 4 which most of them are lower contributed and within-network type. Totally, it seems that global signal regression gives little but different information about the frustration formation. In addition, we did not find any significant differences between frustration formation of lifespan stages as well as without the global signal regression approach.

### Parcellation-based reliability

Many parcellation atlases were developed based on the anatomical and functional attributes of the brain and using different algorithms. In connectivity studies where brain parcellations are used, a question always arises about whether the results are reliable under other atlases (Domhof et al., 2021; Popovych et al., 2021). Actually, we utilized Shen’s parcellation atlas (Shen et al., 2013) and categorized its 268 regions into canonical networks (Yeo et al., 2011) in our analysis. To check the mentioned reliability, we chose Desikan-Killiany-Tourville (DKT) atlas (Klein & Tourville, 2012) as one of the most similar ones to Shen’s parcellation with 101 cortical and subcortical ROIs, although it has different ROI boundaries and lower resolution and was developed based on another algorithm. We also categorized DKT ROIs into canonical networks based on the projections manner that we explained in the method section and Figure S9. Since our analysis was focused on brain elements and considering elemental features of brain atlases are not the same in various atlases, obtaining similar global features and moderate similar local features may be satisfying. Figure S10 shows regional contribution analysis for DKT ROIs and Figure S11 demonstrates that for corresponding functional connections. Both of them do not indicate any significant differences between the contribution map of lifespan stages as the same results were obtained based on Shen’s atlas. Also, subcortical regions and their connections with other brain regions show high contributions to frustration formation that emphasizes the effective role of the subcortex. Most low contributed connections are also within-network type and high contributed connections are almost between subcortical structures and other brain regions. Somatomotor and visual areas also show a low contribution to frustration formation. However, we can see contradictory observations compared to Shen’s related results. For example, a medial large region of the default mode network shows a high contribution opposite to Shen’s results, and the limbic network has some low contributed regions. The differences may be routed in the dissimilarity of the atlases.

### Neural essence of frustration

Frustration is formed by a combination of positive and negative links (Figure 1). Positive and negative links refer to regional synchronous and regional anti-synchronous coactivations, respectively. We don’t know which mechanism emerges negative links, maybe it’s related to time delay due to axonal propagation where influential subcortical regions that are highly contributed to frustration formation have large axonal wiring (Petkoski et al., 2018; Petkoski & Jirsa, 2019). It’s an open question that needs more investigation, and computational modeling can be helpful to solve it. Positive and negative links conceptually look the same as in-phase and anti-phase synchronizes (Petkoski & Jirsa, 2019; Petkoski & Jirsa, 2020; Petkoski et al., 2016) whereas Petkoski and his colleagues showed that in-phase synchrony (or perfectly aligned in/anti-phase clustering) makes the lowest energy which is similar to a brain signed network that has no negative links and frustrations located in the lowest balance energy. They also found that when giving the distribution of the time-delays in the brain, it is more probable that the brain minimizes the disorders that are in a way with our previous results (Saberi et al., 2021a), and the self-organizing essence of the brain (Dresp-Langley, 2020). So we propose their computational approach as a high-potential way to investigate the mechanism behind negative link and frustration appearances.

### Frustration as a brain network measure

Some graph measures are defined based on the arrangement of graph links (Rubinov & Sporns, 2010; Liu et al., 2017), for example, we can calculate the clustering of a network by counting triangles, and motifs as subgraphs are considered the building blocks of the networks. They are commonly investigated in brain networks and extracted from connected-disconnected graphs where either we have a link between two nodes or we don’t have any while frustration is defined in a signed network where presented links have two states of positive and negative and gives information on the system disordering to us.

### Disregarding threshold on connections

Thresholding is a common way of making a brain network (Garrison et al., 2015; Andellini et al., 2015; Wang et al., 2021; Theis et al., 2021). To provide a brain signed network, we claim that although a functional brain network has a low number of anti-synchronous coactivations (Figure S6), the impact of positive and negative connections are not the same, so we should not consider the same threshold for both of them and disregarding the thresholding process is a good way. Consequently, a question emerges that maybe most negative links randomly appear. To answer it we checked the randomness of the contribution pattern (Figure S2-S3). Since we found that they are not random we can conclude that disregarding the threshold is not vulnerable and negative connections appearance make sense.

## Conclusion

In summary, many brain elements play an active role in the frustration formation of the brain network. Although many other elements are less contributed to frustration formation. We identified both of them at the level of the node, connection, and network. The subcortical areas and hippocampus are the most influential region for frustration formation. Matured visual regions have less propensity to get involved in frustrations. Ventral attention to default mode connections are frustrating, especially in adulthood. Regional and connectional contribution patterns are not random. The frustrations are mainly formed between three distinct networks. There is no robust difference between the contribution pattern of brain elements in frustration formation between lifespan stages. The Study of network frustration can reveal the mechanisms behind neural alteration and brain disfunction. Localization of the frustrations also provides the possibility of brain network reorganization.

## Method

### Neuroimage data

We collected functional and anatomical T1 images from two public repositories of ABIDE (Di Martino et al., 2014) and Southwest (Wei et al., 2018). Southwest database contains early adulthood to late adulthood subjects and ABIDE has childhood to early adulthood subjects. We selected all healthy subjects whose functional repetition times were equal to 2 seconds (most frequent repetition times). Selected subjects aged from 6 to 80 (mean: 31.31, sd: 19.78) and 44 percent were female. We classified subjects into 5 lifespan stages of childhood (age: 6–12), adolescence (age: 12–18), early adulthood (age: 18–40), middle adulthood (age: 40–65), and late adulthood (age: greater than 65) according to Erikson’s stages (Sharleen, 2013). After preprocessing, we also excluded subjects whose images could not pass the quality check. Table S6 represents the demography of finalized 793 subjects based on the stage and neuroimages site. Table S7 also describes site-specific imaging protocols. We should mention that the neuroimaging procedures were carried out in compliance with the Declaration of Helsinki. All adult subjects and parents (legal guardians) of subjects under the age of 18 provided informed consent before starting the procedure. Neuroimages are collected in several sites and the acquisition protocols were approved by their licensing committees including the Research Ethics Committee of the Brain Imaging Center of Southwest University, Institutional Review Boards of the New York University School of Medicine, The Institutional Review Board of San Diego State University, the Institutional Review Board of University of Michigan, Yale University Institutional Review Board, Ethics Commission of ETH Zurich, Georgetown University Institutional Review Board, Hospital of Trinity College, and The Institutional Review Board of University of Utah School of Medicine.

### Preprocessing functional images

We employed FSL (Jenkinson et al., 2012) and AFNI (Cox, 1996) to preprocess images. At first, we extracted the brain tissue from the T1 image then segmented it into gray matter (GM), white matter (WM), and cerebral spinal fluid (CSF). Then we removed the first five volumes of the functional image to assure magnetic stability and then performed slice timing correction. After that, we registered volumes of the functional image to the extracted brain of the T1 image using the least square optimization with three translational and three rotational variables. Then we conducted spatial smoothing on registered volumes using a Gaussian kernel (FWHM = 5mm). In the following, we interpolated spiking outliers of every voxel’s time series and applied a bandpass filtering (0.01–0.09 Hz) to them to exclude non-relevant information. We also regressed out three translational and three rotational confounds of motions as well as WM and CSF signals from the time series of every voxel. Finally, we normalized volumes of functional images to MNI152 standard space (2×2×2 mm^3^) by optimization of twelve variables including three translational, three rotational, three scaling, and three shearing variables. We did not regress out the global signal from functional images for analysis of main manuscripts although I brought a version of the results that we regressed out global signals from images in supporting information. In the end, we inspected the quality of pre-processing. So we excluded subjects whose images had low extraction and registration quality and those with movement parameters greater than one voxel size. It’s beneficial to mention that we had used the procedure in other studies (Saberi et al., 2021a; 2021b; Sadeghi et al., 2017; Soheili-Nezhad et al., 2020).

### Regional brain activations

We used MATLAB software to extract 268 regional activity patterns from every preprocessed functional image based on Shen’s atlas (Shen et al., 2013). Although all functional images were acquired with the same repetition time, imaging sites had different acquisition times, volume numbers, and regional time points. The shortest acquisition time belongs to Georgetown University site of ABIDE with 147 volumes. Accordingly, we chose 147 first timepoints of all regional time series. So we obtained 268 time-series with 147 time-points as activity patterns for each subject. Since all repetition times were equal, adjusted activity patterns corresponding to an equal acquisition time. The equality of the number of time points and equality of acquisition times matter in connectivity matrix formation and help to improve the validity of the comparisons.

### Frustration formation

We constructed a connectivity matrix for each subject according to Pearson’s correlation of pairs of regional time series. We only considered signs of correlation coefficients to obtain the adjacency matrix of the subject’s signed network corresponding to positive and negative links. Then we identified the triadic frustrations of the subject’s network. After identifying frustrations, we counted the number of triadic frustrations that every element contributed to in its formation. We measured the contribution value for every nodal, connectional, network-based, and hemisphere elements. We explained the way to estimate the expected contribution value of the elements in the result section.

### Parcellation atlas projection

As we wanted to study the contribution of the canonical networks in frustration formation, we needed to know which Shen’s ROIs belong to which networks. So we calculated the Dice coefficient between any pair of Yeo’s networks and Shen’s ROIs as follows (Meyers et al., 2019; Dice, 1945):

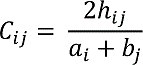

where *a_i_* and *b_j_* correspond to the total voxels in region *i* of Shen’s atlas and the total voxels in network *j* of Yeo’s atlas, respectively. *h_ij_* also denotes the total number of overlapped voxels between them. So we obtained a matrix that each cell indicates a Dice coefficient for a pair of ROI and canonical network (Figure S5). According to the values, we estimated every ROI belongs to which network based on the largest coefficient. Since Yeo only parcellated the cerebral cortex, we grouped subcortical ROIs into three networks of subcortical structures, brainstem, and cerebellum based on their anatomical information. Figure 2 shows the result of ROI categorization. We also attached labeled ROIs in supporting information. In the same way, we also categorized regions of DKT atlases (Figure S9) and provided them to check reliability of results under changing parcellation.

### Entropy calculation

Entropy is a measure of system randomness. When a system has some possible microstates with identical probabilities, Shannon’s entropy determines randomness of variable *X* as follows:

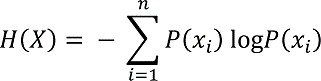

Where summation applies on all states and *P(x_i_)* denotes the probability of the state *x_i_*. The entropy is maximum in the case of equality of states probabilities. We wanted to know whether the appearance of significant nodes and connections in Figure 3 and Figure 4 are random or not. So we calculated Shannon’s entropy for actual patterns and compared them with the entropy of 1000 shuffled patterns. In this regard, we measured the entropy of regional patterns as follows. Firstly, we considered canonical networks as states, counted the number of significant regions located in each network, normalized counted numbers to bin size (total number of ROIs located in networks), then divided the outputs by the total number of significant ROIs. In this way, we provided probabilities of states and calculated Shannon’s entropy for the apparent pattern of significant brain regions (Figure S2). As similar, we measured Shannon’s entropy for the pattern of significant brain connections but we regarded connections between networks as states (Figure S3).

### Statistical analysis

In this study, we had two types of comparison for each element: the two-group paired comparison between actual and null contribution values, and multiple-group comparison between contribution values of lifespan stages. We used Wilcoxon matched-pairs signed-rank test for the first one and the Kruskal-Wallis test for the last one. We utilized non-parametric statistical analysis because contribution values were not distributed normally. We performed multiple comparison corrections on p-values using False Discovery Rate (FDR) and considered effect size to improve the validity of the statistical analysis. We also used two different algorithms for calculating effect sizes of paired between-group and multiple-group analyses with different thresholds (Table 2). Also, we calculated the p-values of entropy analysis based on the null hypothesis of “entropy of actual pattern is larger than entropies of shuffled patterns”. We should mention that we carried out all statistical analyses in R software (RC Team, 2013; Mangiafico & Mangiafico, 2017; Kassambara, 2020; Whitcher et al., 2013). We provided Figure 1 using “draw.io” and statistical figures and brain maps by the advance of “BrainNet Viewer” (Xia et al., 2013) and some other R packages (Nakazawa, 2019; Wickham et al., 2016).

**Table 2.**
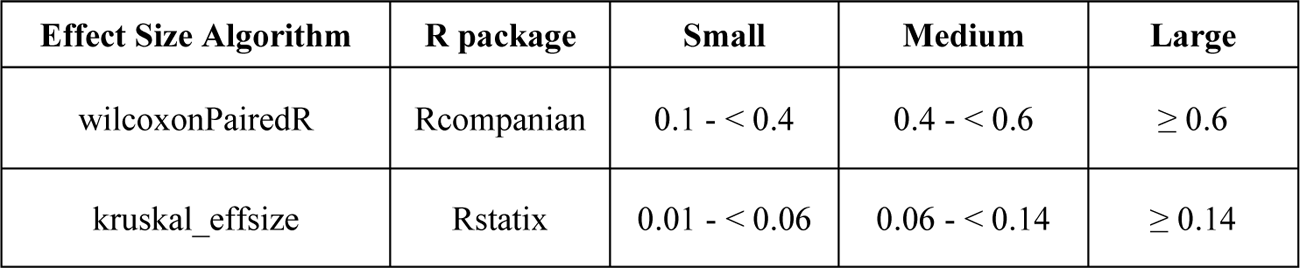
Effect size thresholds and corresponding algorithms.

In addition, we shared all of the information and codes on “https://github.com/majidsaberi/BrainNetFrustration” so everyone can publicly access, replicate, and develop our research.

## Supporting information

Shen2Yeo Labels

Shen2Yeo Projection

Connectional Contribution Statistics

Regional Contribution Statistics

Shen2Yeo Projection - Network 1

Shen2Yeo Projection - Network 2

Shen2Yeo Projection - Network 3

Shen2Yeo Projection - Network 4

Shen2Yeo Projection - Network 5

Shen2Yeo Projection - Network 6

Shen2Yeo Projection - Network 7

Shen2Yeo Projection - Network 8

Shen2Yeo Projection - Network 9

Shen2Yeo Projection - Network 10

## Contributions

M.S designed the study. R.K, A.K, B.M, and G.J advised the research. M.S analyzed the data and wrote the manuscript. R.K, A.K, B.M, and G.J revised the manuscript.

### Corresponding author

Majid Saberi

---

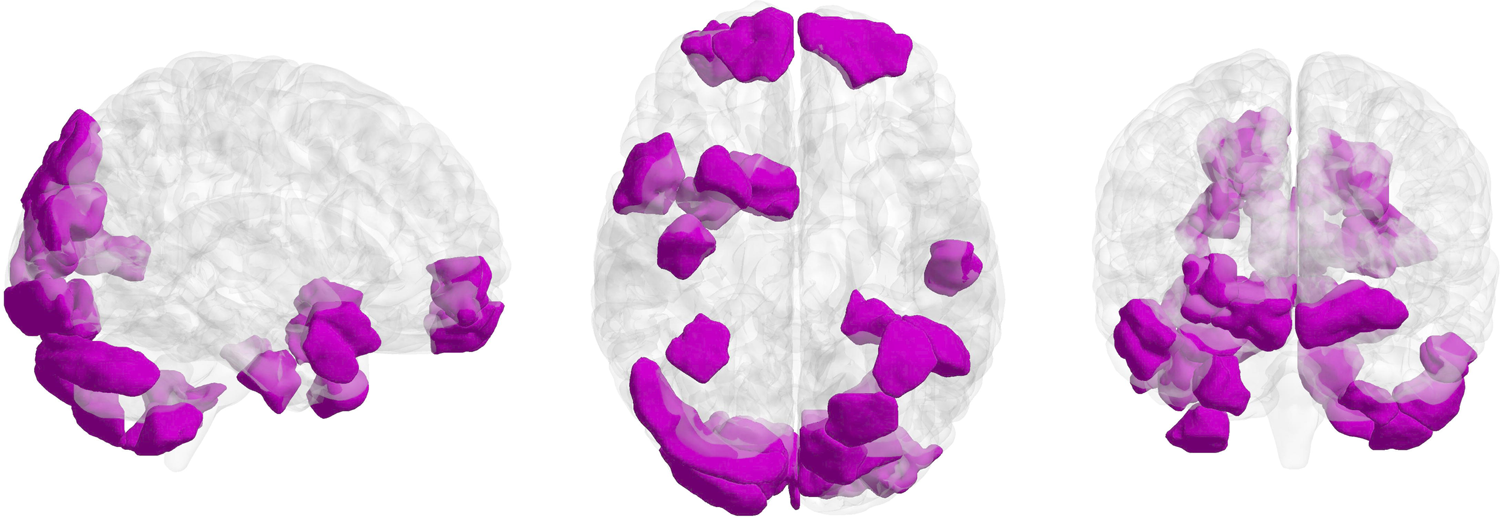

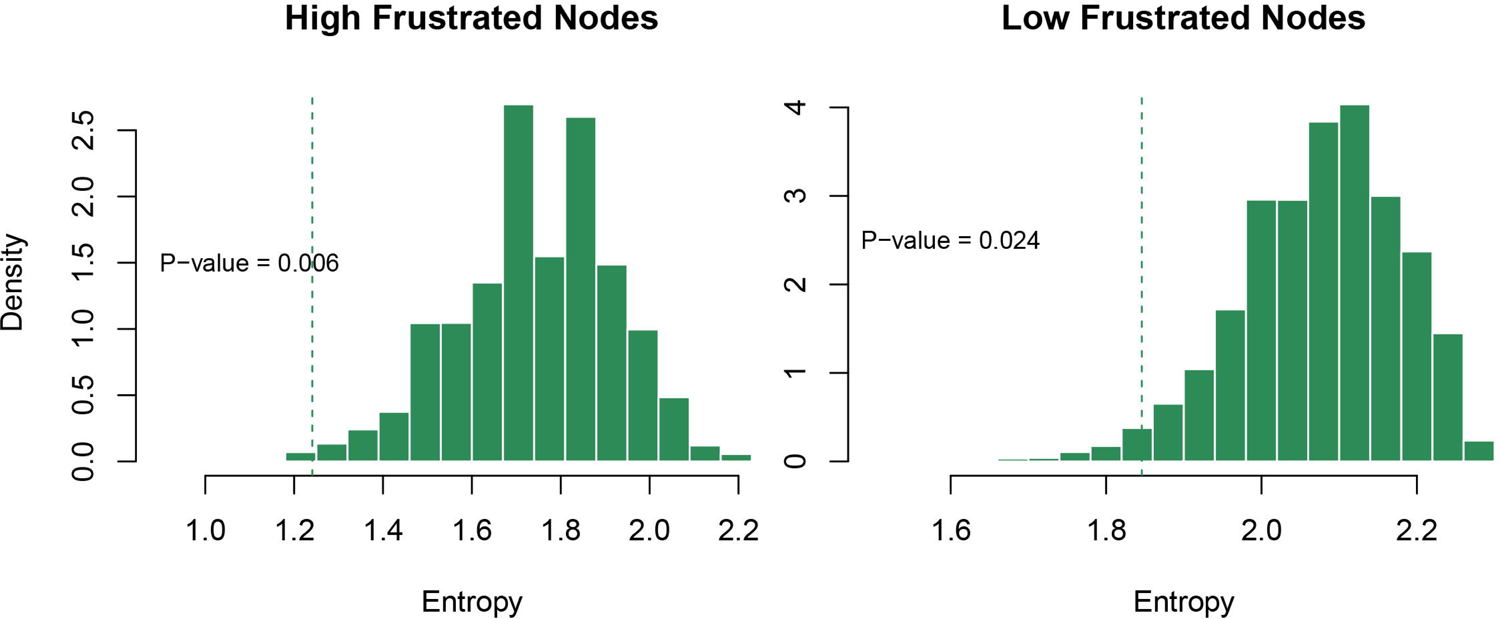

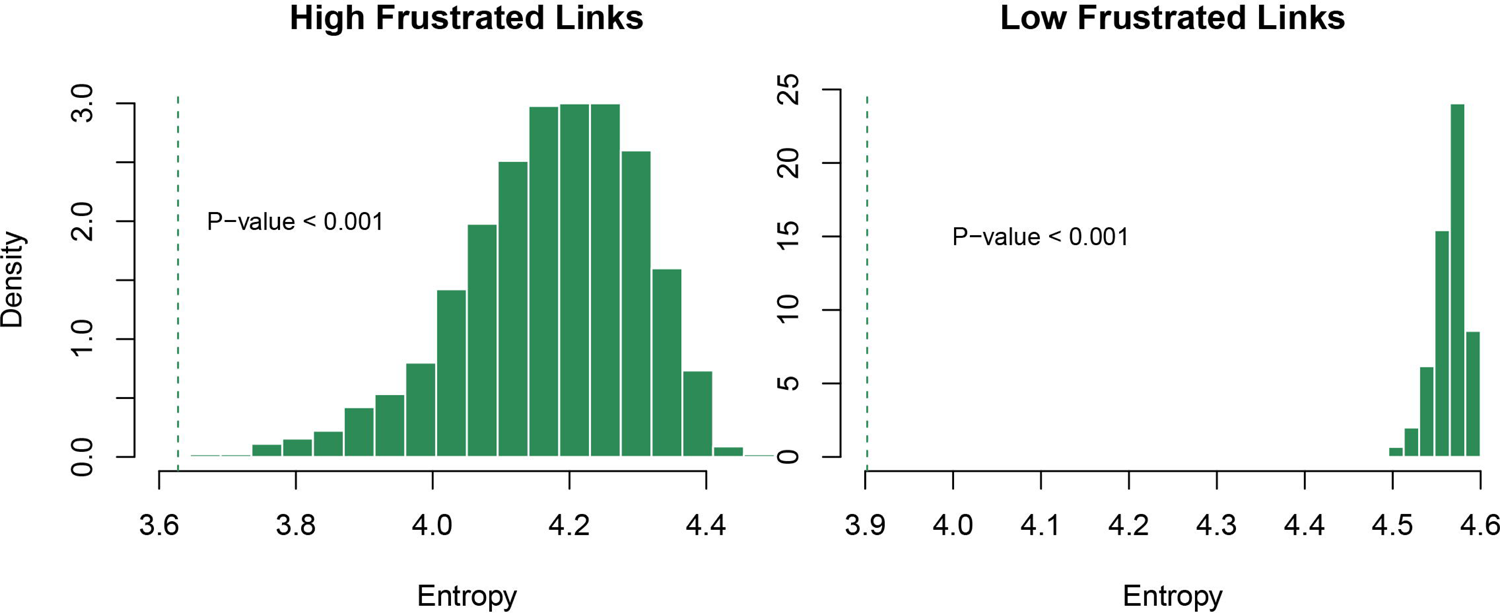

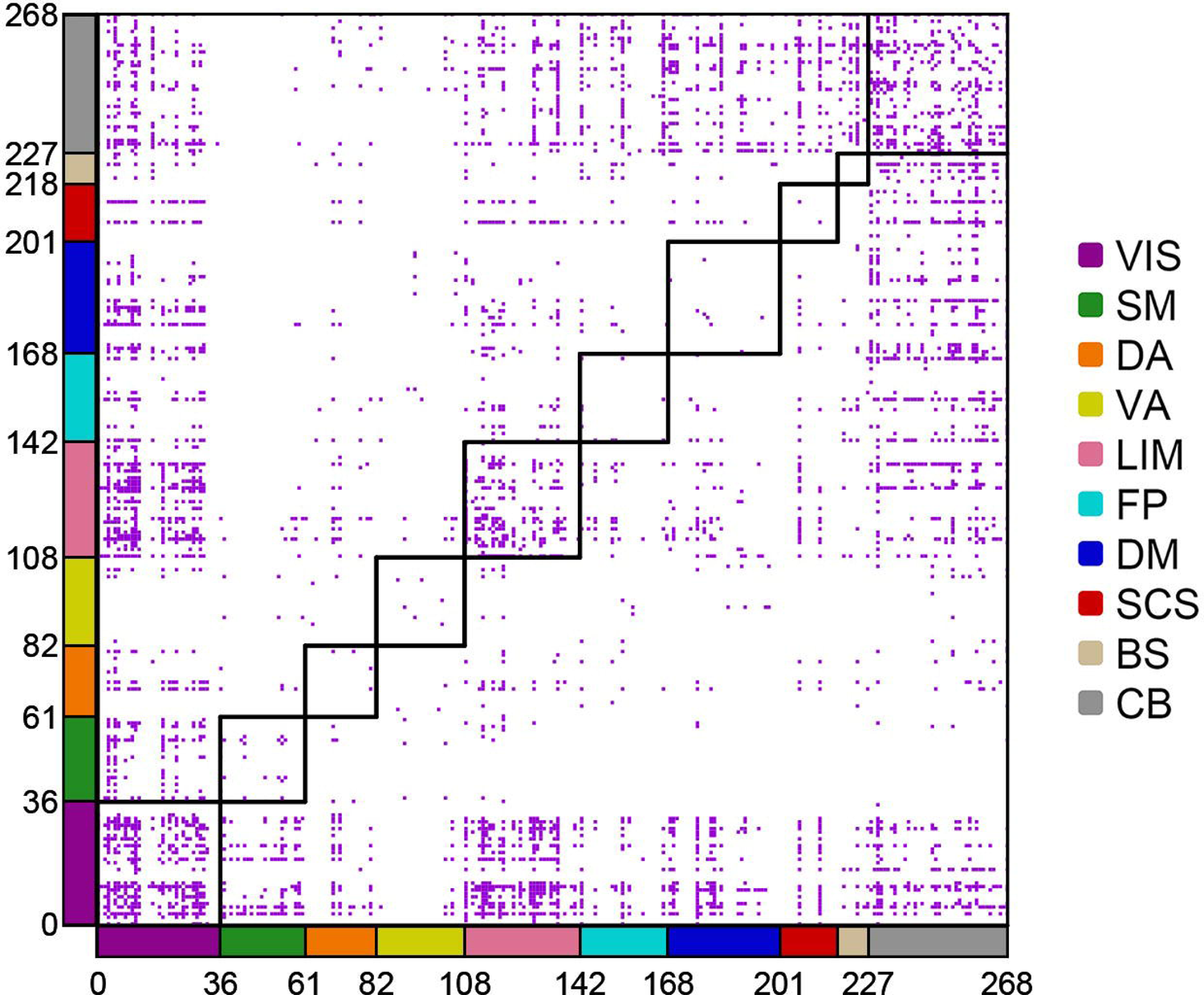

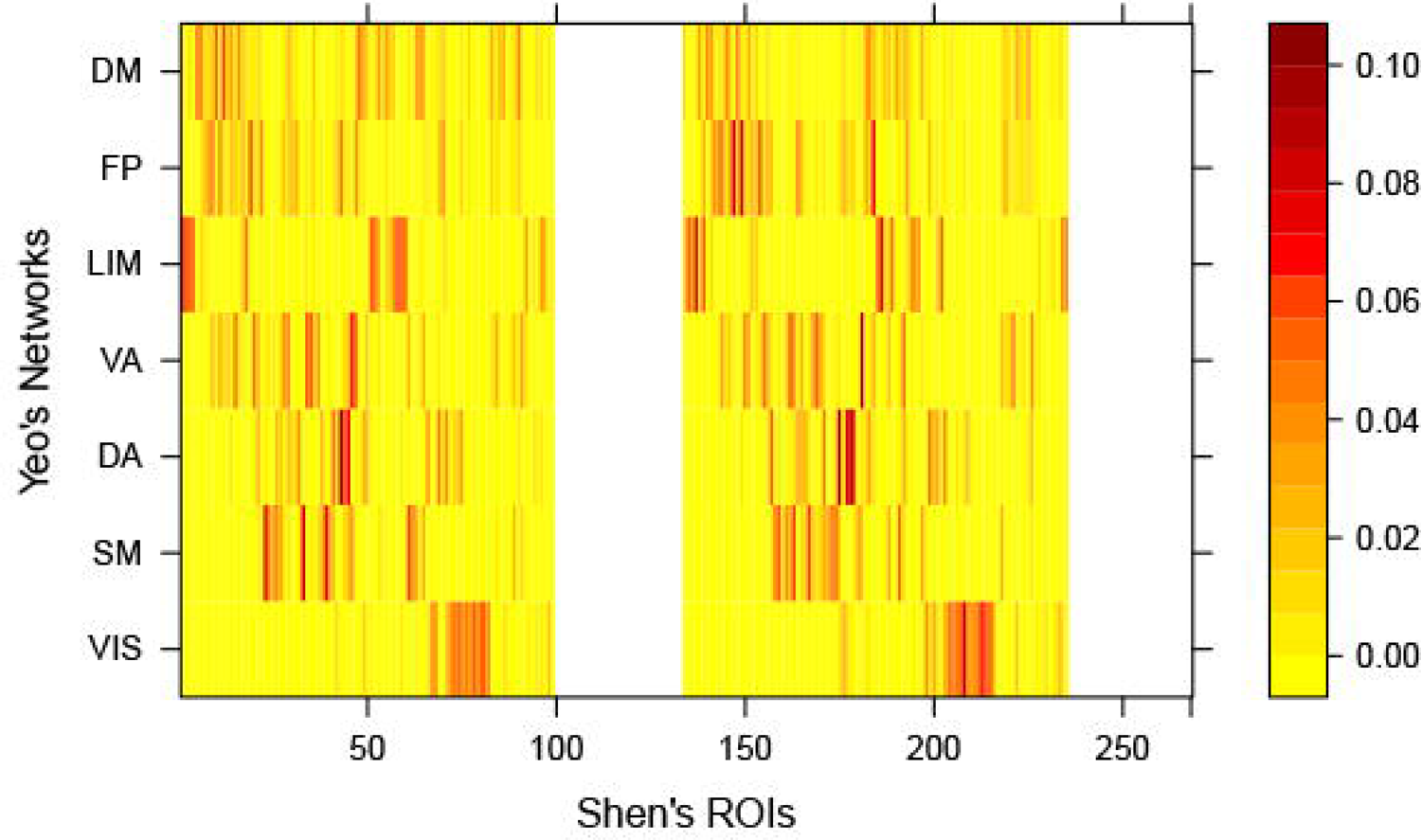

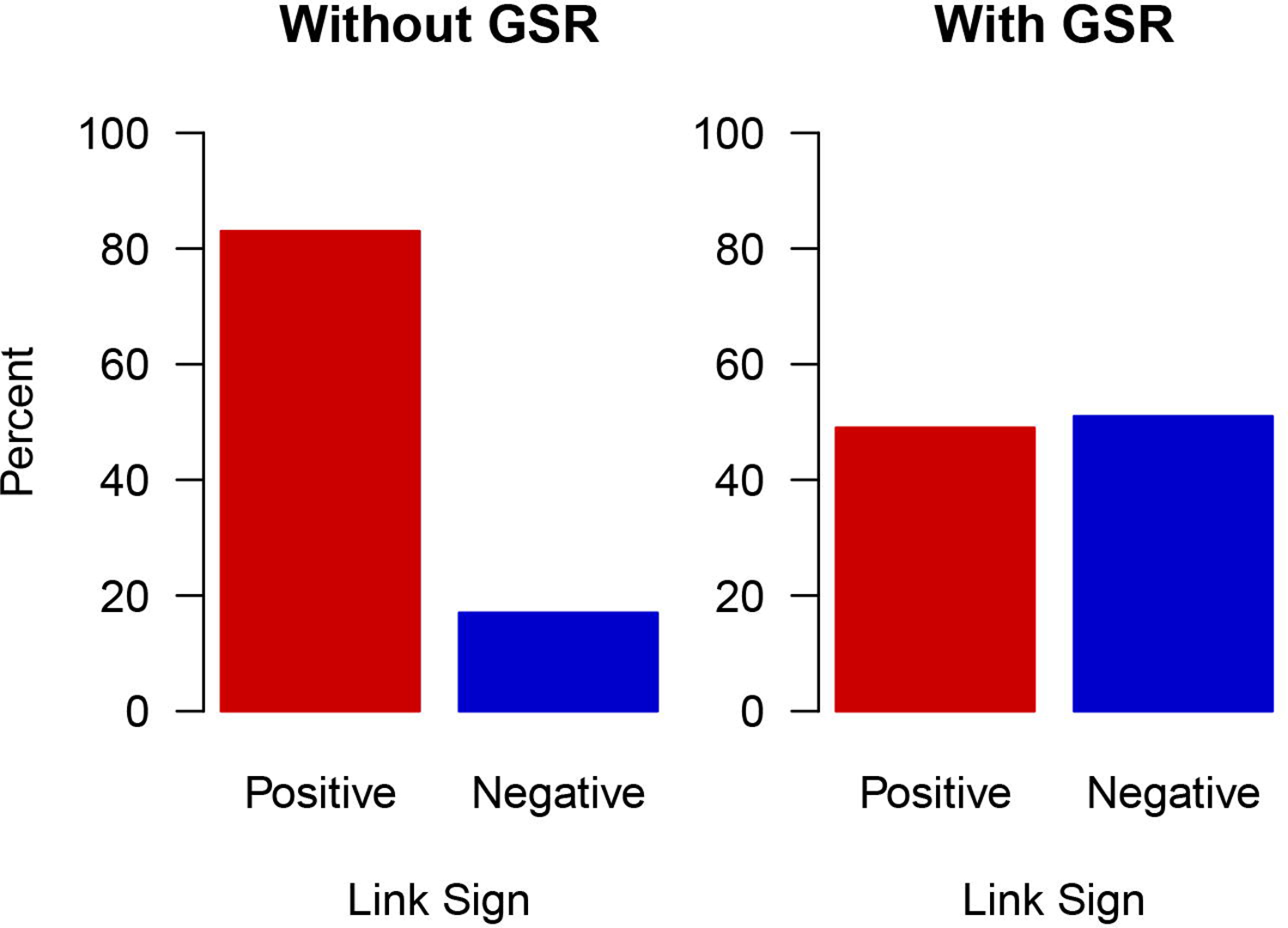

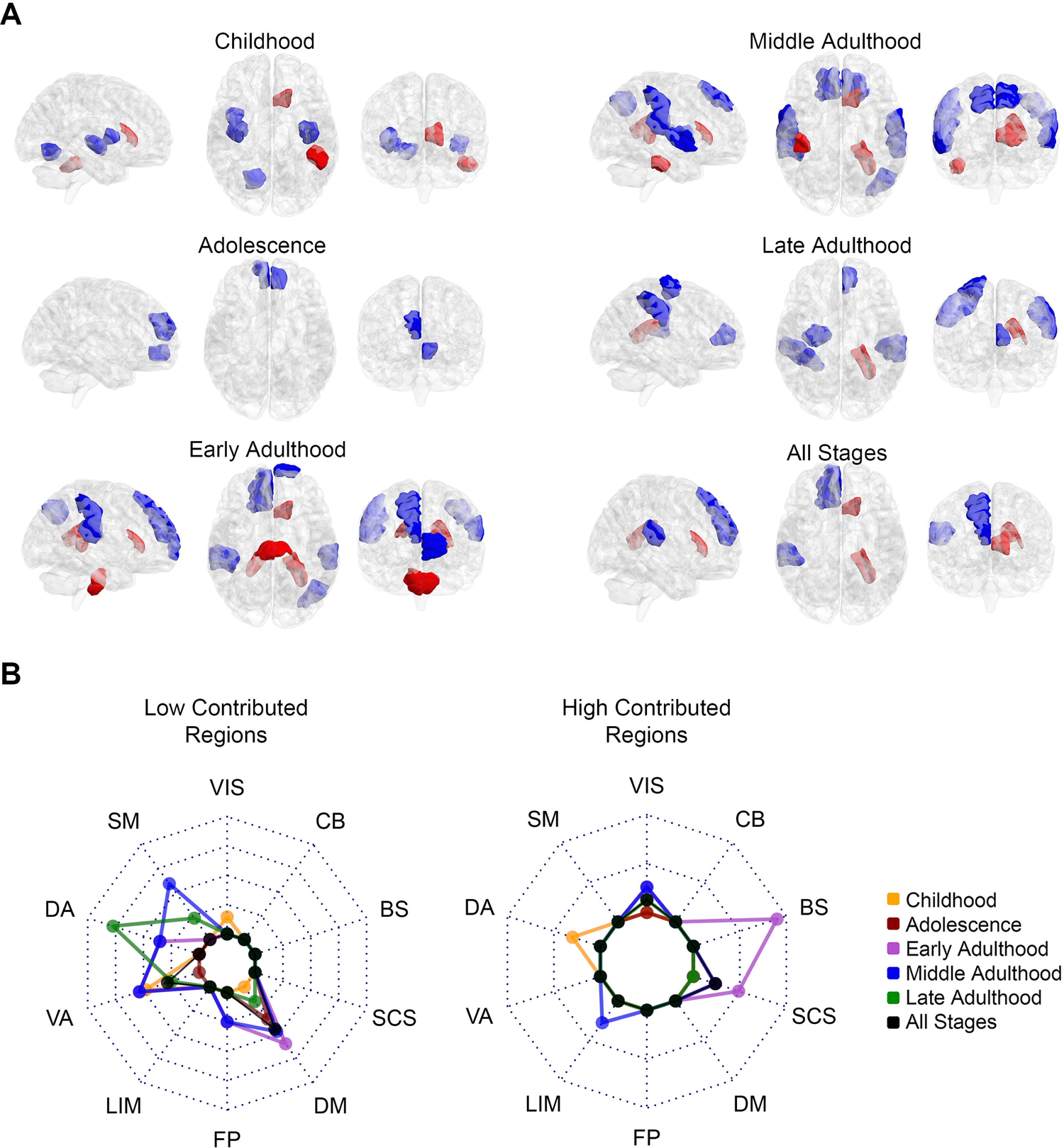

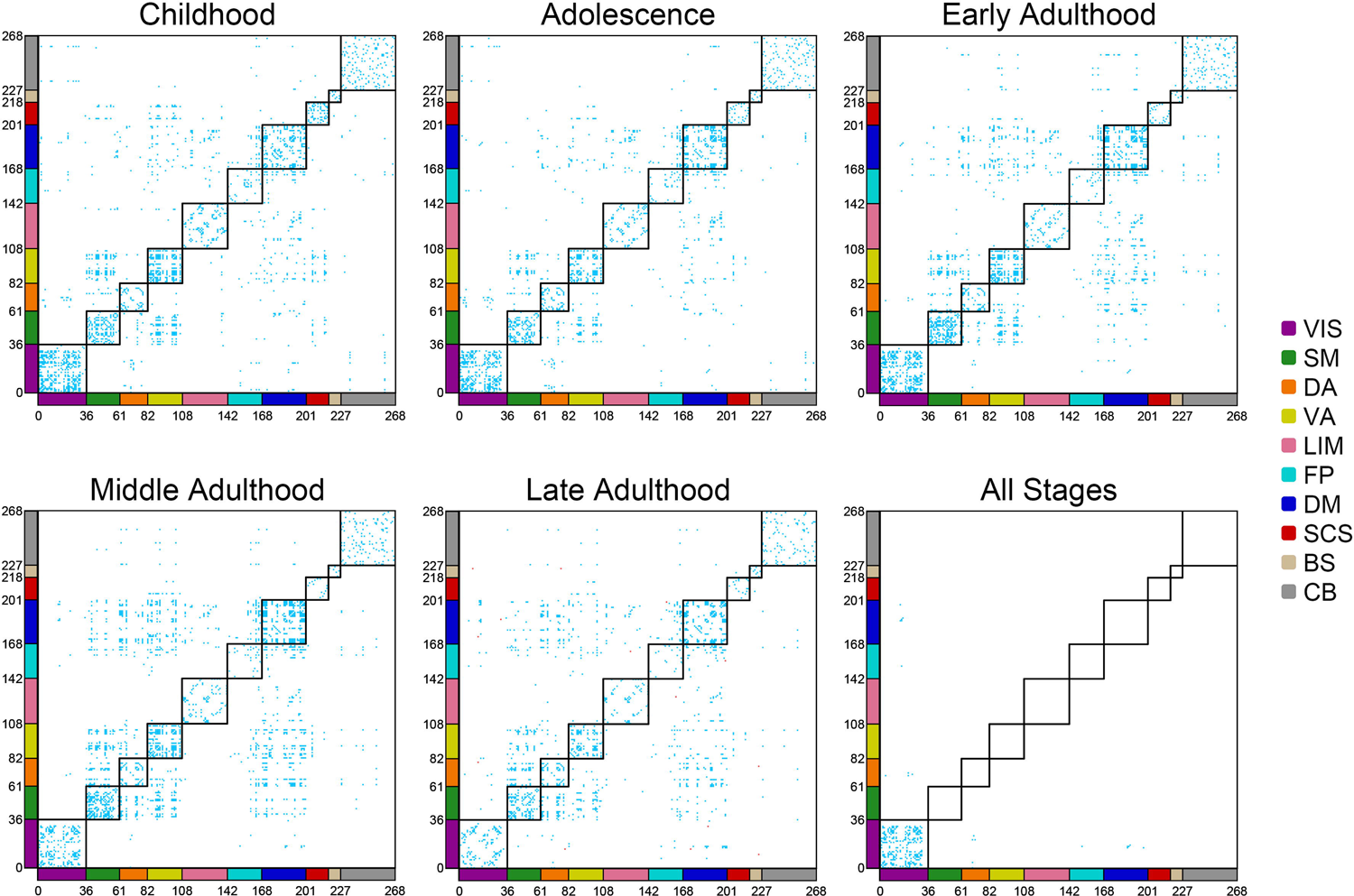

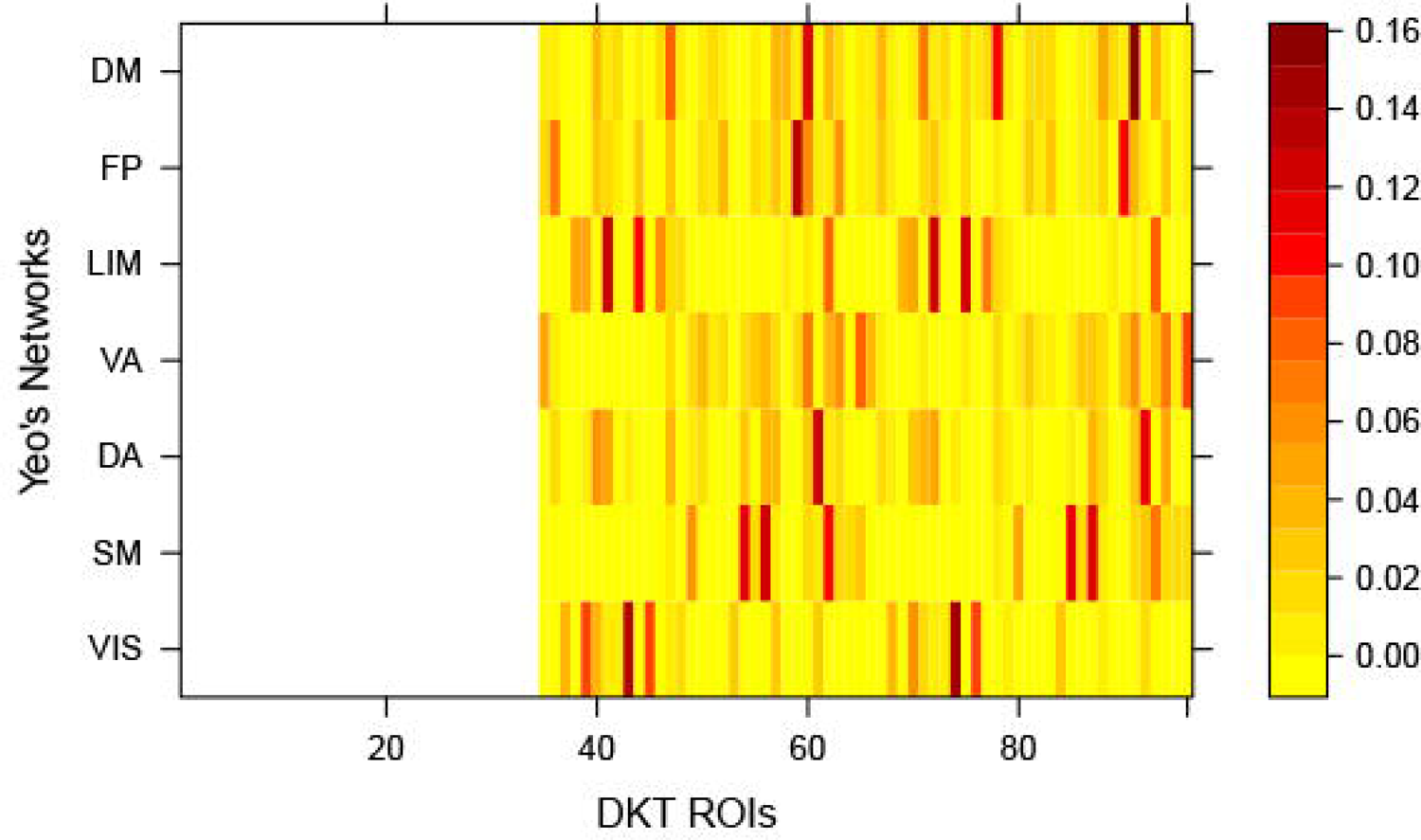

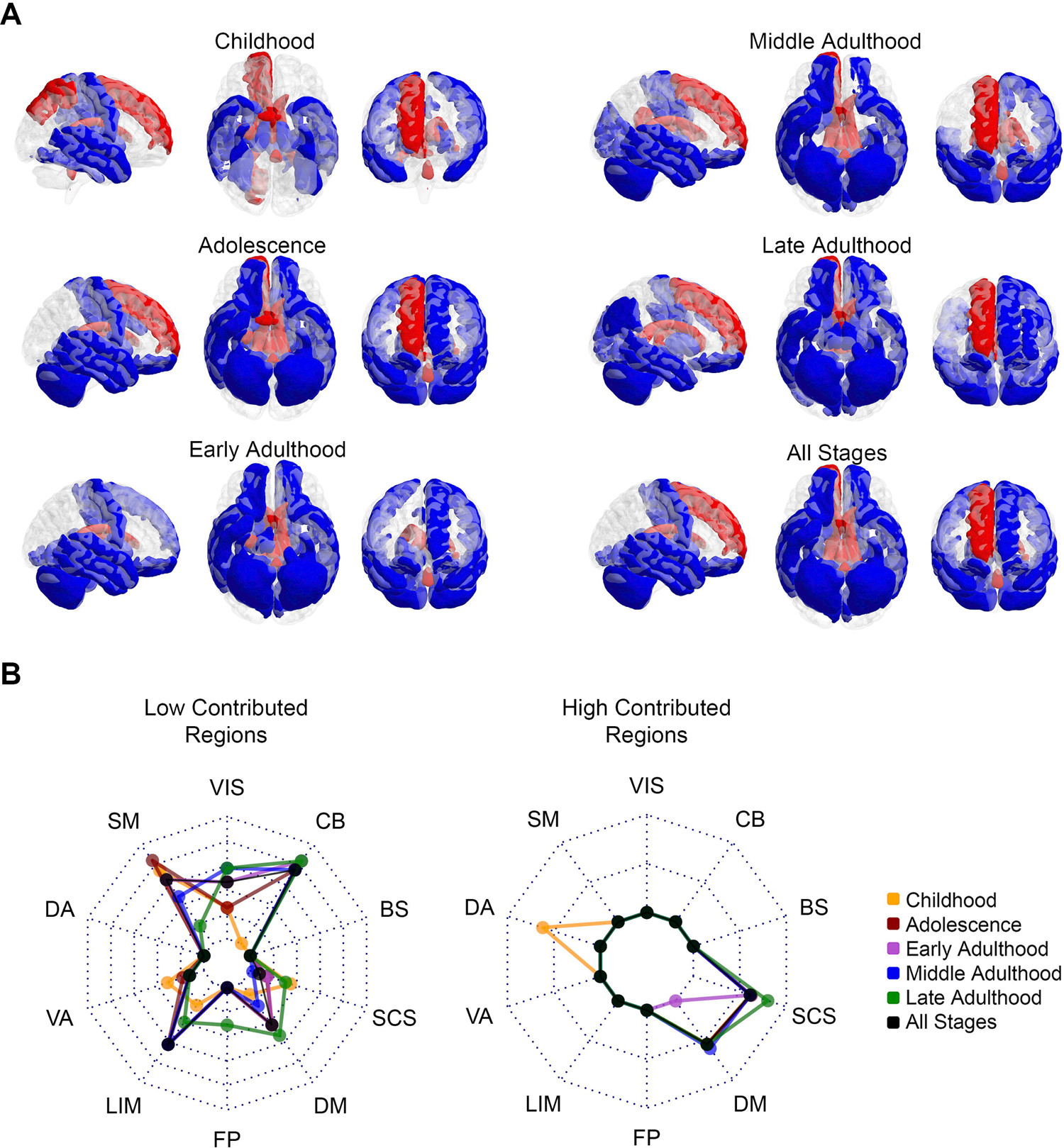

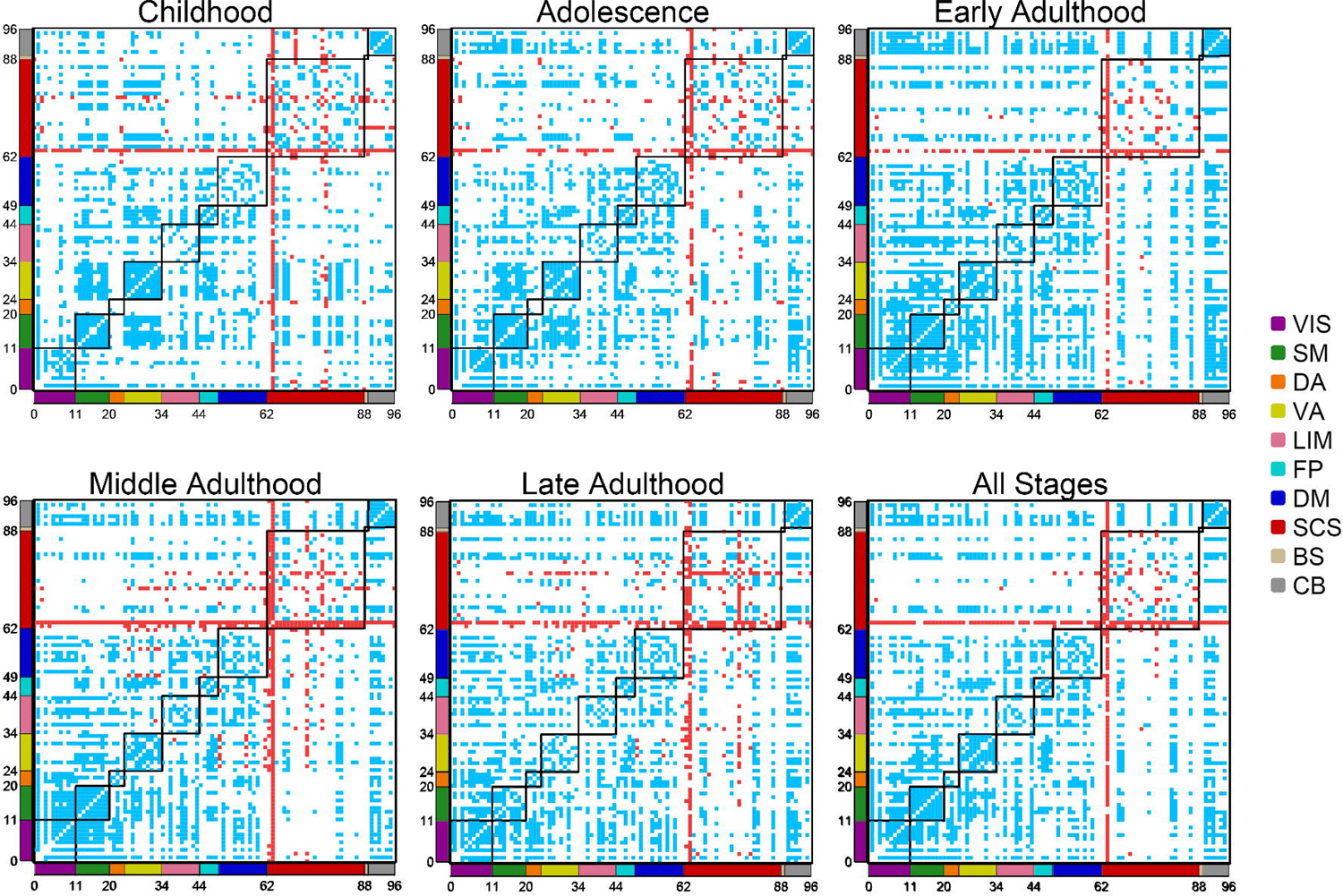

## References

1. Anand, K. S., & Dhikav, V. (2012). Hippocampus in health and disease: An overview. Annals of Indian Academy of Neurology, 15(4), 239.

2. Anchuri, P., & Magdon-Ismail, M. (2012, August). Communities and balance in signed networks: A spectral approach. In 2012 IEEE/ACM International Conference on Advances in Social Networks Analysis and Mining (pp. 235-242). IEEE.

3. Andellini, M., Cannatà, V., Gazzellini, S., Bernardi, B., & Napolitano, A. (2015). Test-retest reliability of graph metrics of resting state MRI functional brain networks: A review. Journal of neuroscience methods, 253, 183–192.

4. Antal, T., Krapivsky, P. L., & Redner, S. (2005). Dynamics of social balance on networks. Physical Review E, 72(3), 036121.

5. Aquino, K. M., Fulcher, B. D., Parkes, L., Sabaroedin, K., & Fornito, A. (2020). Identifying and removing widespread signal deflections from fMRI data: Rethinking the global signal regression problem. Neuroimage, 212, 116614.

6. Aref, S., & Wilson, M. C. (2019). Balance and frustration in signed networks. Journal of Complex Networks, 7(2), 163–189.

7. Arnold Anteraper, S., Triantafyllou, C., Sawyer, A. T., Hofmann, S. G., Gabrieli, J. D., & Whitfield-Gabrieli, S. (2014). Hyper-connectivity of subcortical resting-state networks in social anxiety disorder. Brain connectivity, 4(2), 81–90.

8. Awad, M. Z., Vaden, R. J., Irwin, Z. T., Gonzalez, C. L., Black, S., Nakhmani, A., … & Walker, H. C. (2021). Subcortical short-term plasticity elicited by deep brain stimulation. Annals of clinical and translational neurology, 8(5), 1010–1023.

9. Bassett, D. S., & Bullmore, E. T. (2009). Human brain networks in health and disease. Current opinion in neurology, 22(4), 340.

10. Bassett, D. S., & Sporns, O. (2017). Network neuroscience. Nature neuroscience, 20(3), 353–364.

11. Cartwright, D., & Harary, F. (1956). Structural balance: a generalization of Heider’s theory. Psychological review, 63(5), 277.

12. Cerliani, L., Mennes, M., Thomas, R. M., Di Martino, A., Thioux, M., & Keysers, C. (2015). Increased functional connectivity between subcortical and cortical resting-state networks in autism spectrum disorder. JAMA psychiatry, 72(8), 767–777.

13. Chandrasekaran, B., Skoe, E., & Kraus, N. (2014). An integrative model of subcortical auditory plasticity. Brain topography, 27(4), 539–552.

14. Chen, Y. L., Tu, P. C., Huang, T. H., Bai, Y. M., Su, T. P., Chen, M. H., & Wu, Y. T. (2020). Using minimal-redundant and maximal-relevant whole-brain functional connectivity to classify bipolar disorder. Frontiers in neuroscience, 1098.

15. Colle, H., Colle, D., Noens, B., Dhaen, B., Alessi, G., Muller, P., … & van der Linden, C. (2021). Subcortical Stimulation with Tip of Ultrasound Aspirator. Journal of Neurological Surgery Part A: Central European Neurosurgery, 82(06), 581–584.

16. Cox, R. W. (1996). AFNI: software for analysis and visualization of functional magnetic resonance neuroimages. Computers and Biomedical research, 29(3), 162–173.

17. Derr, T., Aggarwal, C., & Tang, J. (2018, October). Signed network modeling based on structural balance theory. In Proceedings of the 27th ACM international conference on information and knowledge management (pp. 557-566).

18. Di Martino, A., Yan, C. G., Li, Q., Denio, E., Castellanos, F. X., Alaerts, K., … & Milham, M. P. (2014). The autism brain imaging data exchange: towards a large-scale evaluation of the intrinsic brain architecture in autism. Molecular psychiatry, 19(6), 659–667.

19. Dice, L. R. (1945). Measures of the amount of ecologic association between species. Ecology, 26(3), 297–302.

20. Domakonda, M. J., He, X., Lee, S., Cyr, M., & Marsh, R. (2019). Increased functional connectivity between ventral attention and default mode networks in adolescents with bulimia nervosa. Journal of the American Academy of Child & Adolescent Psychiatry, 58(2), 232–241.

21. Domhof, J. W., Jung, K., Eickhoff, S. B., & Popovych, O. V. (2021). Parcellation-induced variation of empirical and simulated brain connectomes at group and subject levels. Network Neuroscience, 5(3), 798–830.

22. Dresp-Langley, B. (2020). Seven properties of self-organization in the human brain. Big Data and Cognitive Computing, 4(2), 10.

23. Duffau, H. (2009). Does post-lesional subcortical plasticity exist in the human brain?. Neuroscience Research, 65(2), 131–135.

24. Duménieu, M., Marquèze-Pouey, B., Russier, M., & Debanne, D. (2021). Mechanisms of Plasticity in Subcortical Visual Areas. Cells, 10(11), 3162.

25. Estrada, E. (2019). Rethinking structural balance in signed social networks. Discrete Applied Mathematics, 268, 70–90.

26. Facchetti, G., Iacono, G., & Altafini, C. (2011). Computing global structural balance in large-scale signed social networks. Proceedings of the National Academy of Sciences, 108(52), 20953–20958.

27. Finn, E. S., Shen, X., Scheinost, D., Rosenberg, M. D., Huang, J., Chun, M. M., … & Constable, R. T. (2015). Functional connectome fingerprinting: identifying individuals using patterns of brain connectivity. Nature neuroscience, 18(11), 1664–1671.

28. Folloni, D., Verhagen, L., Mars, R. B., Fouragnan, E., Constans, C., Aubry, J. F., … & Sallet, J. (2019). Manipulation of subcortical and deep cortical activity in the primate brain using transcranial focused ultrasound stimulation. Neuron, 101(6), 1109–1116.

29. Fox, M. D., Zhang, D., Snyder, A. Z., & Raichle, M. E. (2009). The global signal and observed anticorrelated resting state brain networks. Journal of neurophysiology, 101(6), 3270–3283.

30. Garrison, K. A., Scheinost, D., Finn, E. S., Shen, X., & Constable, R. T. (2015). The (in) stability of functional brain network measures across thresholds. Neuroimage, 118, 651–661.

31. Goremychkin, E. A., Osborn, R., Rainford, B. D., Macaluso, R. T., Adroja, D. T., & Koza, M. (2008). Spin-glass order induced by dynamic frustration. Nature Physics, 4(10), 766–770.

32. Heider, F. (1946). Attitudes and cognitive organization. The Journal of psychology, 21(1), 107–112.

33. Hoogman, M., Bralten, J., Hibar, D. P., Mennes, M., Zwiers, M. P., Schweren, L. S., … & Franke, B. (2017). Subcortical brain volume differences in participants with attention deficit hyperactivity disorder in children and adults: a cross-sectional mega-analysis. The Lancet Psychiatry, 4(4), 310–319.

34. Jenkinson, M., Beckmann, C. F., Behrens, T. E., Woolrich, M. W., & Smith, S. M. (2012). Fsl. Neuroimage, 62(2), 782–790.

35. Jones, E. G. (2000). Cortical and subcortical contributions to activity-dependent plasticity in primate somatosensory cortex. Annual review of neuroscience, 23(1), 1–37.

36. Kassambara, A. (2020). rstatix: Pipe-friendly framework for basic statistical tests. R package version 0.6.0.

37. Kirkley, A., Cantwell, G. T., & Newman, M. E. (2019). Balance in signed networks. Physical Review E, 99(1), 012320.

38. Klein, A., & Tourville, J. (2012). 101 labeled brain images and a consistent human cortical labeling protocol. Frontiers in neuroscience, 6, 171.

39. Lanciego, J. L., Luquin, N., & Obeso, J. A. (2012). Functional neuroanatomy of the basal ganglia. Cold Spring Harbor perspectives in medicine, 2(12), a009621.

40. Lee, Y. J. G., Kim, S., Kim, N., Choi, J. W., Park, J., Kim, S. J., … & Lee, Y. J. (2018). Changes in subcortical resting-state functional connectivity in patients with psychophysiological insomnia after cognitive–behavioral therapy. NeuroImage: Clinical, 17, 115–123.

41. Liu, D., Wang, S., Gao, Q., Dong, R., Fu, X., Pugh, E., & Hu, J. (2020). Learning a second language in adulthood changes subcortical neural encoding. Neural Plasticity, 2020.

42. Liu, J., Li, M., Pan, Y., Lan, W., Zheng, R., Wu, F. X., & Wang, J. (2017). Complex brain network analysis and its applications to brain disorders: a survey. Complexity, 2017.

43. Liu, T. T., Nalci, A., & Falahpour, M. (2017). The global signal in fMRI: Nuisance or Information?. Neuroimage, 150, 213–229.

44. Mangiafico, S., & Mangiafico, M. S. (2017). Package ‘rcompanion’. Cran Repos, 20, 1–71.

45. McElroy, S. L., Kotwal, R., Keck Jr, P. E., & Akiskal, H. S. (2005). Comorbidity of bipolar and eating disorders: distinct or related disorders with shared dysregulations?. Journal of affective disorders, 86(2-3), 107-127.

46. Meyers, P. E., Arvapalli, G. C., Ramachandran, S. C., Frank, P. F., Lemmer, A. D., Bridgeford, E. W., & Vogelstein, J. T. (2019). Standardizing human brain parcellations. *Biorxiv*, October.

47. Miranda, J. A., Shepard, K. N., McClintock, S. K., & Liu, R. C. (2014). Adult plasticity in the subcortical auditory pathway of the maternal mouse. PLoS One, 9(7), e101630.

48. Mišić, B., & Sporns, O. (2016). From regions to connections and networks: new bridges between brain and behavior. Current opinion in neurobiology, 40, 1–7.

49. Murphy, K., Birn, R. M., Handwerker, D. A., Jones, T. B., & Bandettini, P. A. (2009). The impact of global signal regression on resting state correlations: are anti-correlated networks introduced?. Neuroimage, 44(3), 893–905.

50. Nakazawa, M., & Nakazawa, M. M. (2019). Package ‘fmsb’. *See* https://cran.r-project.org/web/packages/fmsb/fmsb.pdf.

51. Petkoski, S., & Jirsa, V. K. (2019). Transmission time delays organize the brain network synchronization. Philosophical Transactions of the Royal Society A, 377(2153), 20180132.

52. Petkoski, S., & Jirsa, V. K. (2020). Normalizing the brain connectome for communication through synchronization. Network Neuroscience, 1-56.

53. Petkoski, S., Palva, J. M., & Jirsa, V. K. (2018). Phase-lags in large scale brain synchronization: Methodological considerations and in-silico analysis. PLoS computational biology, 14(7), e1006160.

54. Petkoski, S., Spiegler, A., Proix, T., Aram, P., Temprado, J. J., & Jirsa, V. K. (2016). Heterogeneity of time delays determines synchronization of coupled oscillators. Physical Review E, 94(1), 012209.

55. Popovych, O. V., Jung, K., Manos, T., Diaz-Pier, S., Hoffstaedter, F., Schreiber, J., … & Eickhoff, S. B. (2021). Inter-subject and inter-parcellation variability of resting-state whole-brain dynamical modeling. Neuroimage, 236, 118201.

56. Rapoport, A. (1963). Mathematical models of social interaction.

57. RC Team, (2013). R: A language and environment for statistical computing.

58. Rosenberg, M. D., Finn, E. S., Scheinost, D., Papademetris, X., Shen, X., Constable, R. T., & Chun, M. M. (2016). A neuromarker of sustained attention from whole-brain functional connectivity. Nature neuroscience, 19(1), 165–171.

59. Rosenberg-Katz, K., Herman, T., Jacob, Y., Kliper, E., Giladi, N., & Hausdorff, J. M. (2016). Subcortical volumes differ in Parkinson’s disease motor subtypes: new insights into the pathophysiology of disparate symptoms. Frontiers in human neuroscience, 10, 356.

60. Rubinov, M., & Sporns, O. (2010). Complex network measures of brain connectivity: uses and interpretations. Neuroimage, 52(3), 1059–1069.

61. Saberi, M., Khosrowabadi, R., Khatibi, A., Misic, B., & Jafari, G. (2021). Topological impact of negative links on the stability of resting-state brain network. Scientific reports, 11(1), 1–14.

62. Saberi, M., Khosrowabadi, R., Khatibi, A., Misic, B., & Jafari, G. (2021). Requirement to change of functional brain network across the lifespan. PloS one, 16(11), e0260091.

63. Sadeghi, M., Khosrowabadi, R., Bakouie, F., Mahdavi, H., Eslahchi, C., & Pouretemad, H. (2017). Screening of autism based on task-free fmri using graph theoretical approach. Psychiatry Research: Neuroimaging, 263, 48–56.

64. Sampaio-Baptista, C., Khrapitchev, A. A., Foxley, S., Schlagheck, T., Scholz, J., Jbabdi, S., … & Johansen-Berg, H. (2013). Motor skill learning induces changes in white matter microstructure and myelination. Journal of Neuroscience, 33(50), 19499–19503.

65. Schölvinck, M. L., Maier, A., Frank, Q. Y., Duyn, J. H., & Leopold, D. A. (2010). Neural basis of global resting-state fMRI activity. Proceedings of the National Academy of Sciences, 107(22), 10238–10243.

66. Scholz, J., Klein, M. C., Behrens, T. E., & Johansen-Berg, H. (2009). Training induces changes in white-matter architecture. Nature neuroscience, 12(11), 1370–1371.

67. Sharleen, L. K. (2013). Lifespan development. Goodheart-Wilcox Publisher.

68. Shen, X., Tokoglu, F., Papademetris, X., & Constable, R. T. (2013). Groupwise whole-brain parcellation from resting-state fMRI data for network node identification. Neuroimage, 82, 403–415.

69. Sheykhali, S., Darooneh, A. H., & Jafari, G. R. (2020). Partial balance in social networks with stubborn links. Physica A: Statistical Mechanics and its Applications, 548, 123882.

70. Shi, S., Xu, A. G., Rui, Y. Y., Zhang, X., Romanski, L. M., Gothard, K. M., & Roe, A. W. (2021). Infrared neural stimulation with 7T fMRI: A rapid in vivo method for mapping cortical connections of primate amygdala. NeuroImage, 231, 117818.

71. Shi, Y., Ikrar, T., Olivas, N. D., & Xu, X. (2014). Bidirectional global spontaneous network activity precedes the canonical unidirectional circuit organization in the developing hippocampus. Journal of Comparative Neurology, 522(9), 2191–2208.

72. Siu, C. R., & Murphy, K. M. (2018). The development of human visual cortex and clinical implications. Eye and brain, 10, 25.

73. Soheili-Nezhad, S., Jahanshad, N., Guelfi, S., Khosrowabadi, R., Saykin, A. J., Thompson, P. M., … & Zarei, M. (2020). Imaging genomics discovery of a new risk variant for Alzheimer’s disease in the postsynaptic SHARPIN gene. Human brain mapping, 41(13), 3737–3748.

74. Tang, J., Chang, Y., Aggarwal, C., & Liu, H. (2016). A survey of signed network mining in social media. ACM Computing Surveys (CSUR), 49(3), 1–37.

75. Theis, N., Rubin, J., Cape, J., Iyengar, S., Gur, R. E., Gur, R. C., … & Prasad, K. M. (2021). Evaluating Network Threshold Selection for Structural and Functional Brain Connectomes. bioRxiv.

76. Toulouse, G. (1987). Theory of the frustration effect in spin glasses: I. Spin Glass Theory and Beyond: An Introduction to the Replica Method and Its Applications, 9, 99.

77. Vannimenus, J., & Toulouse, G. (1977). Theory of the frustration effect. II. Ising spins on a square lattice. Journal of Physics C: Solid State Physics, 10(18), L537.

78. Villain, J., Bidaux, R., Carton, J. P., & Conte, R. (1980). Order as an effect of disorder. Journal de Physique, 41(11), 1263–1272.

79. Walsh, B. T., Roose, S. P., Glassman, A. H., Gladis, M., & Sadik, C. (1985). Bulimia and depression. Psychosomatic Medicine.

80. Wang, Z., Jie, B., Feng, C., Wang, T., Bian, W., Ding, T. X., … & Liu, M. (2021). Distribution-guided Network Thresholding for Functional Connectivity Analysis in fMRI-based Brain Disorder Identification. IEEE journal of biomedical and health informatics.

81. Wei, D., Zhuang, K., Ai, L., Chen, Q., Yang, W., Liu, W., … & Qiu, J. (2018). Structural and functional brain scans from the cross-sectional Southwest University adult lifespan dataset. Scientific data, 5(1), 1–10.

82. Whitcher, B., Schmid, V., Thornton, A., Whitcher, M. B., & Suggests, X. M. L. (2013). Package ‘oro. nifti’.

83. Wickham, H., Chang, W., & Wickham, M. H. (2016). Package ‘ggplot2’. Create elegant data visualisations using the grammar of graphics. Version, 2(1), 1–189.

84. Winkler, M., & Reichardt, J. (2013). Motifs in triadic random graphs based on Steiner triple systems. Physical Review E, 88(2), 022805.

85. Xia, M., Wang, J., & He, Y. (2013). BrainNet Viewer: a network visualization tool for human brain connectomics. PloS one, 8(7), e68910.

86. Yang, B., Cheung, W., & Liu, J. (2007). Community mining from signed social networks. IEEE transactions on knowledge and data engineering, 19(10), 1333–1348.

87. Yang, S. H., Smola, A. J., Long, B., Zha, H., & Chang, Y. (2012, August). Friend or frenemy? Predicting signed ties in social networks. In Proceedings of the 35th international ACM SIGIR conference on Research and development in information retrieval (pp. 555-564).

88. Yeo, B. T., Krienen, F. M., Sepulcre, J., Sabuncu, M. R., Lashkari, D., Hollinshead, M., … & Buckner, R. L. (2011). The organization of the human cerebral cortex estimated by intrinsic functional connectivity. Journal of neurophysiology.

89. Zhao, H., Zhang, J., Lyu, M., Bachus, S., Tokiwa, Y., Gegenwart, P., … & Sun, P. (2019). Quantum-critical phase from frustrated magnetism in a strongly correlated metal. Nature Physics, 15(12), 1261–1266.

