## Supporting Information for "Pattern of frustration formation in the functional brain network"

#### Title:

### Figures:

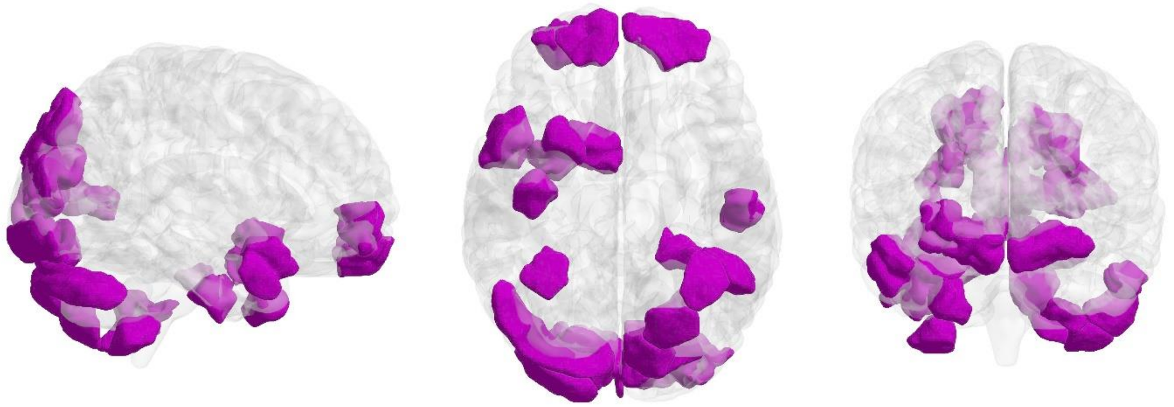

**Figure S1. Between-stage comparison of regional contribution in frustration formation.** Violet regions denote ROIs that have between-stage differences with corrected p-value lower than 0.05 and medium effect sizes.

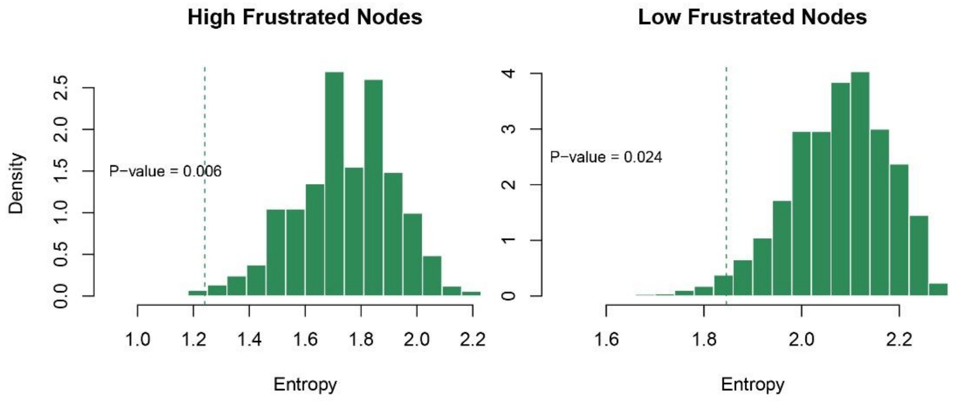


**Figure S2. Comparison of actual regional pattern's entropies with shuffled regional pattern's entropies.** The left and right subfigures refer to regions with significantly higher contribution and regions with the significantly lower contribution in frustration formation. Histograms show shuffled pattern's entropies and vertical dashed lines denote to the value of the actual pattern's entropies shown in Figure 3.

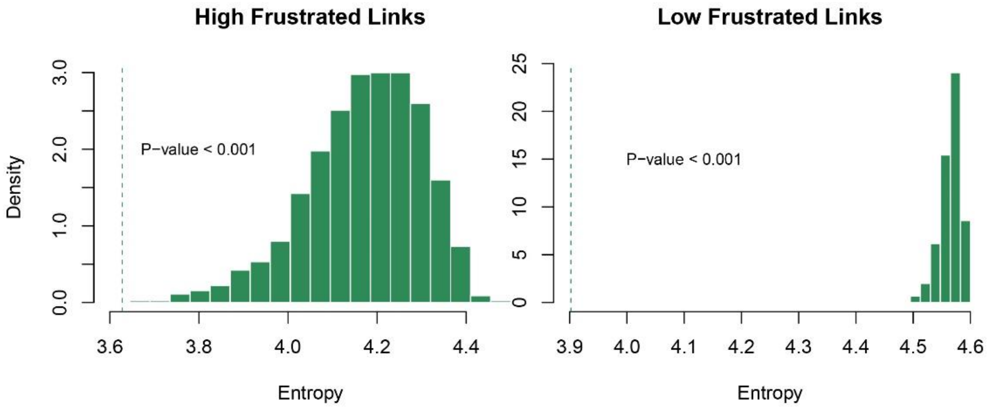


**Figure S3. Comparison of actual connectional pattern's entropies with shuffled connectional pattern's entropies.** The left and right subfigures refer to connections with significantly higher contribution and connections with the significantly lower contribution in frustration formation. Histograms show shuffled pattern's entropies and vertical dashed lines denote the value of the actual pattern's entropies shown in Figure 4.

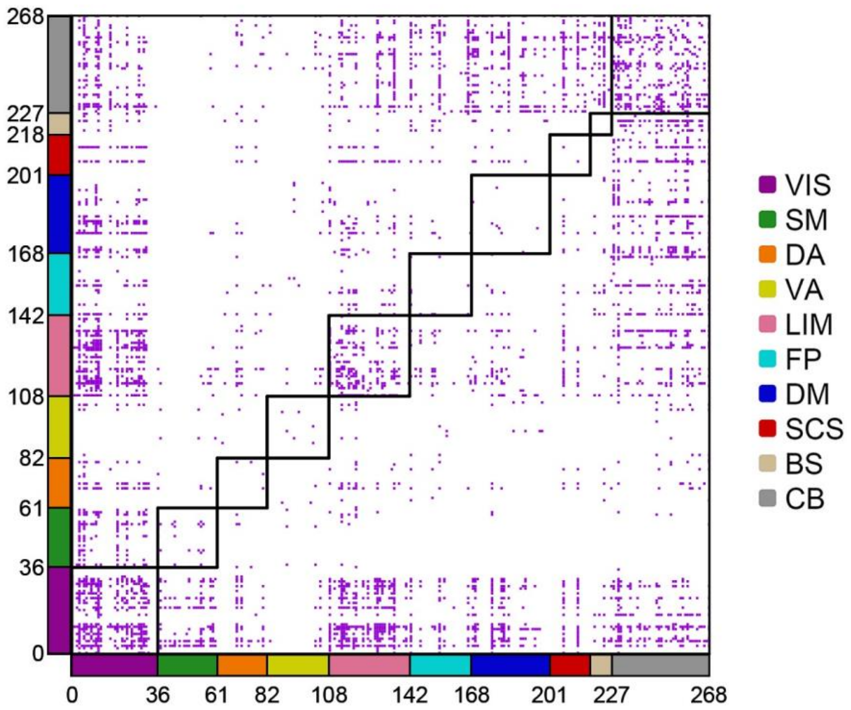


**Figure S4. Between-stage comparison of connectional contribution in frustration formation.** Shen's 268 ROIs are categorized into 10 networks of Fig. 2 by axes coloring. Cells also represent functional connections between every two ROIs. Violet cells denote connections that have between-stage differences with corrected p-value lower than 0.05 and medium effect sizes. Black squares discriminate between network connections. Abbreviation; VIS: visual, SM: somatomotor, DA: dorsal attention, VA: ventral attention, LIM: limbic, FP: frontoparietal, DM: default mode, SCS: subcortical structures, BS: brainstem, CB: cerebellum.

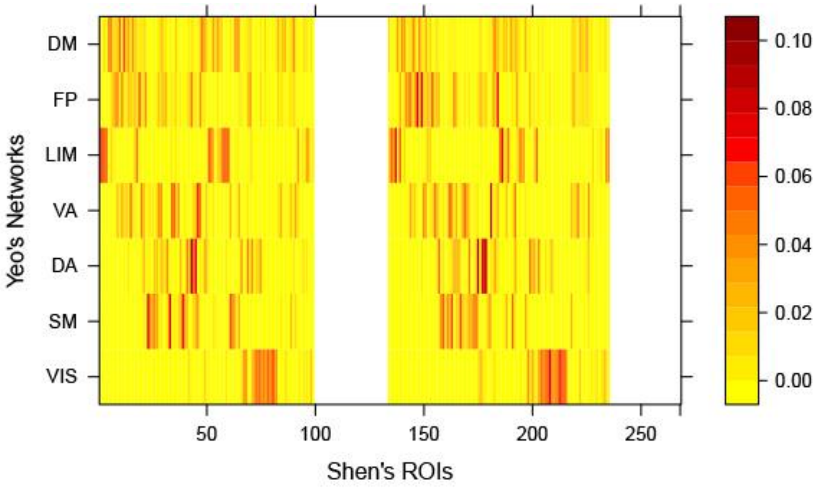


**Figure S5. Projection of Shen’s ROIs into Yeo’s canonical networks.** Colorbar denotes the Dice coefficient for each pair of ROI and canonical networks. Yeo’s atlas is a cortical parcellation so subcortical ROIs can not project into it, corresponding null cells. Abbreviation; VIS: visual, SM: somatomotor, DA: dorsal attention, VA: ventral attention, LIM: limbic, FP: frontoparietal, DM: default mode.

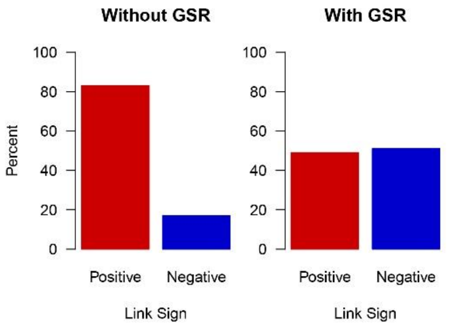


**Figure S6. Percentages of positive and negative links before and after Global Signal Regression (GSR).**

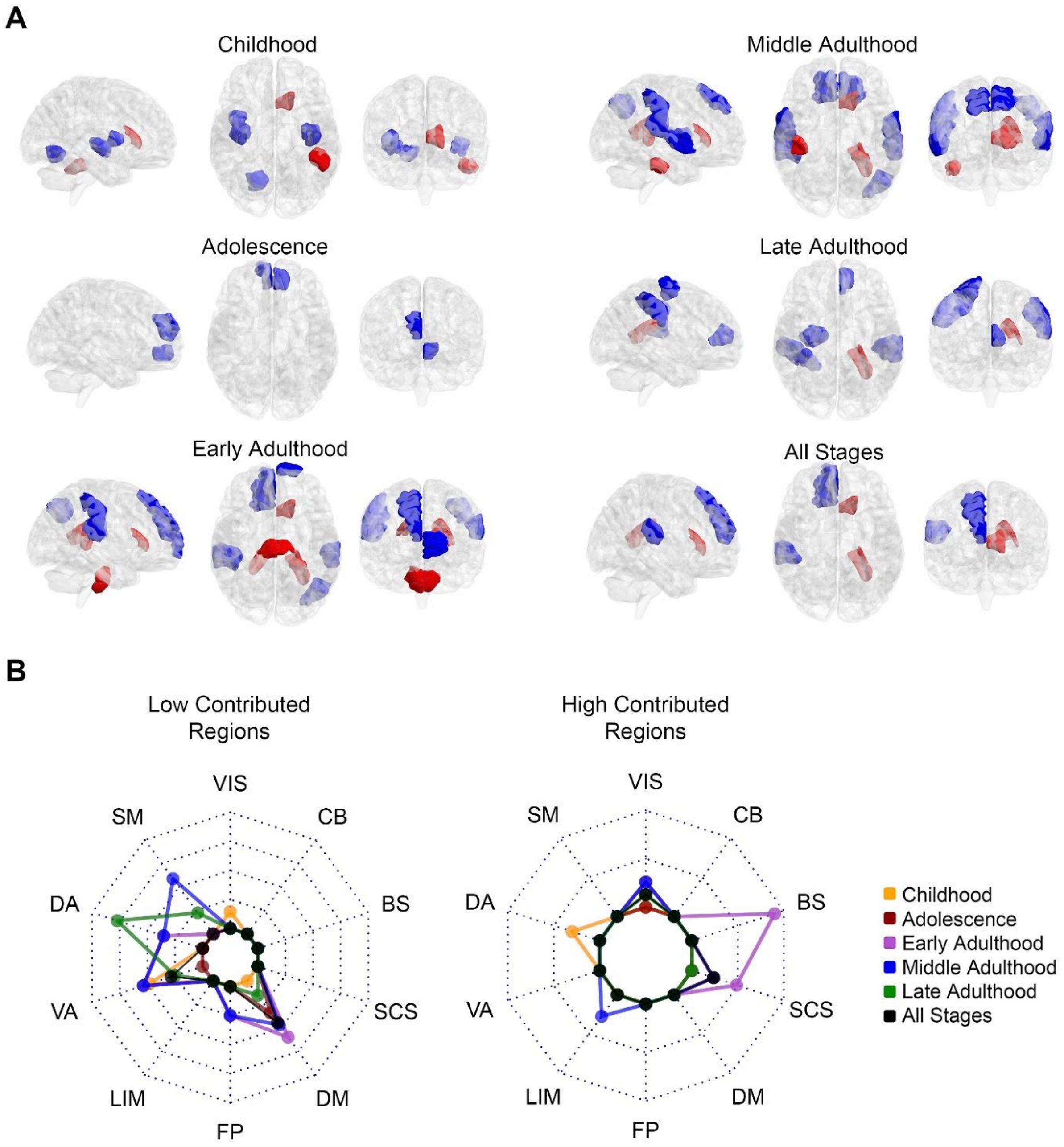


**Figure S7. The pattern of regional contribution in frustration formation when we added global signal regression to preprocessing.** Red-colored and blue-colored areas of the brain maps indicate Shen's regions with significantly greater and significantly lower contributions in frustration formation. Brain maps demonstrate the patterns for various lifespan stages and all stages in three representational planes of sagittal, axial, and coronal. Abbreviation; VIS: visual, SM: somatomotor, DA: dorsal attention, VA: ventral attention, LIM: limbic, FP: frontoparietal, DM: default mode, SCS: subcortical structures, BS: brainstem, CB: cerebellum.

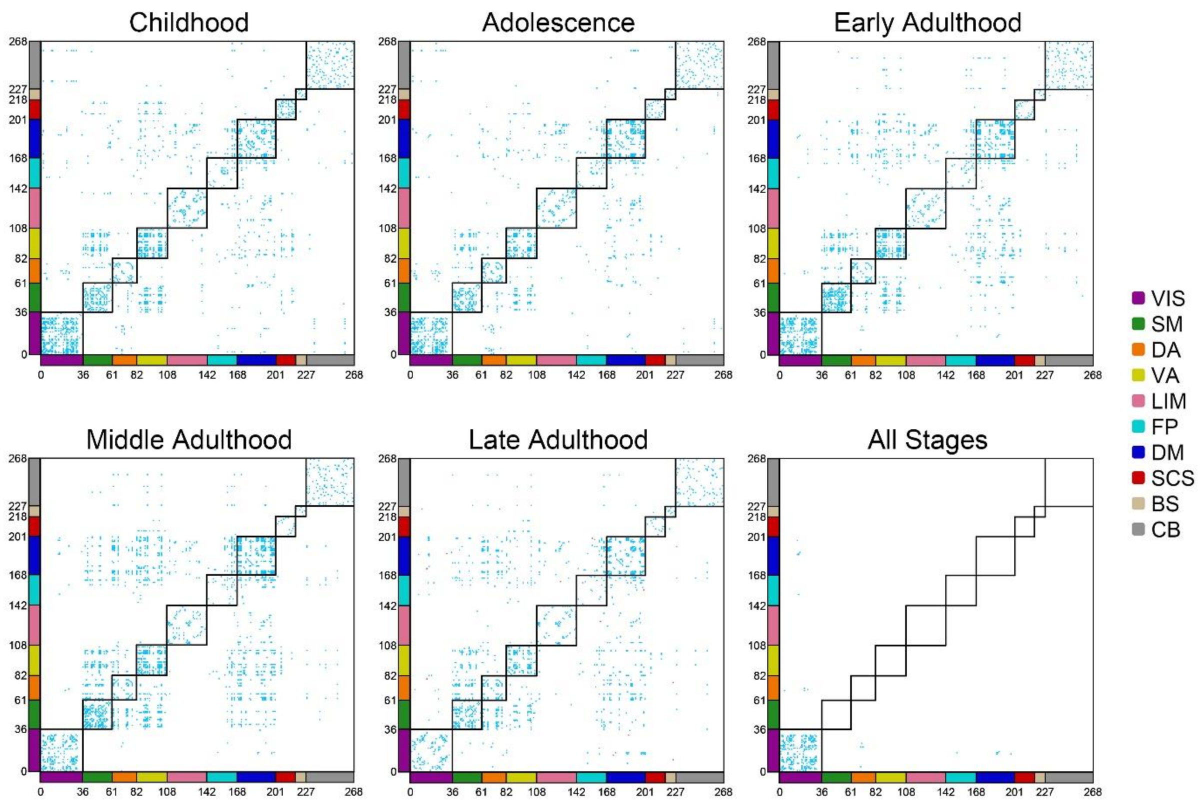


**Figure S8 - Contribution maps of functional connections in frustration formation when we added global signal regression to preprocessing.** The first five maps demonstrate the maps related to lifespan stages and the last one corresponds without considering the stage. Shen's 268 ROIs are categorized into 10 networks of Figure 2 by axes coloring. Cells also represent functional connections between every two ROIs. Red cells display those connections with a significantly greater contribution to frustration formation and blue cells indicate those connections with significantly lower involvement. Black squares discriminate between network connections. Abbreviation; VIS: visual, SM: somatomotor, DA: dorsal attention, VA: ventral attention, LIM: limbic, FP: frontoparietal, DM: default mode, SCS: subcortical structures, BS: brainstem, CB: cerebellum.

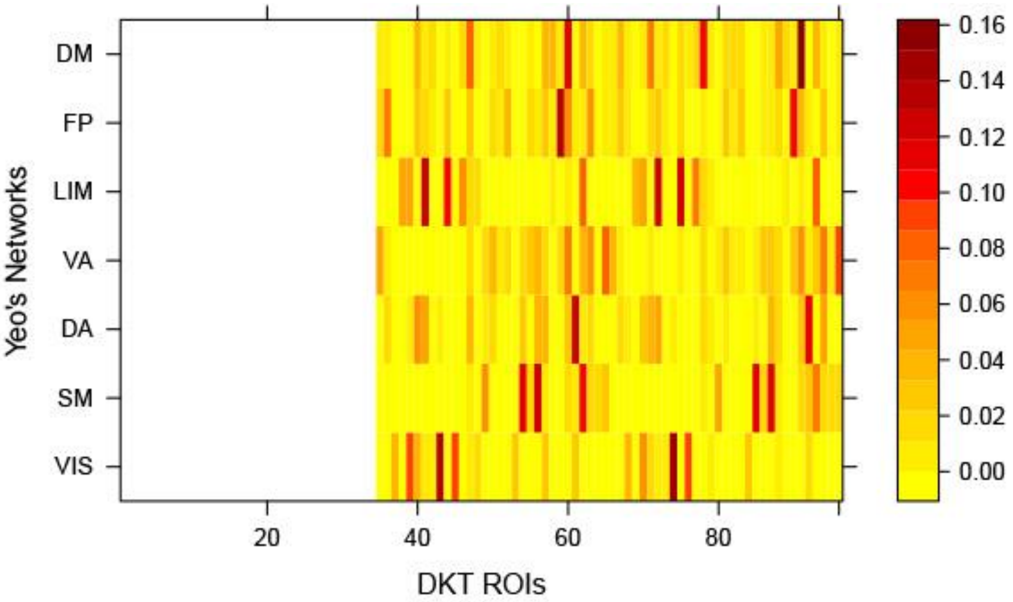


**Figure S9. Projection of DKT atlas ROIs into Yeo's canonical networks.** Colorbar denotes the Dice coefficient for each pair of ROI and canonical networks. Yeo's atlas is a cortical parcellation so subcortical ROIs can not project into it, corresponding null cells. Abbreviation; VIS: visual, SM: somatomotor, DA: dorsal attention, VA: ventral attention, LIM: limbic, FP: frontoparietal, DM: default mode.

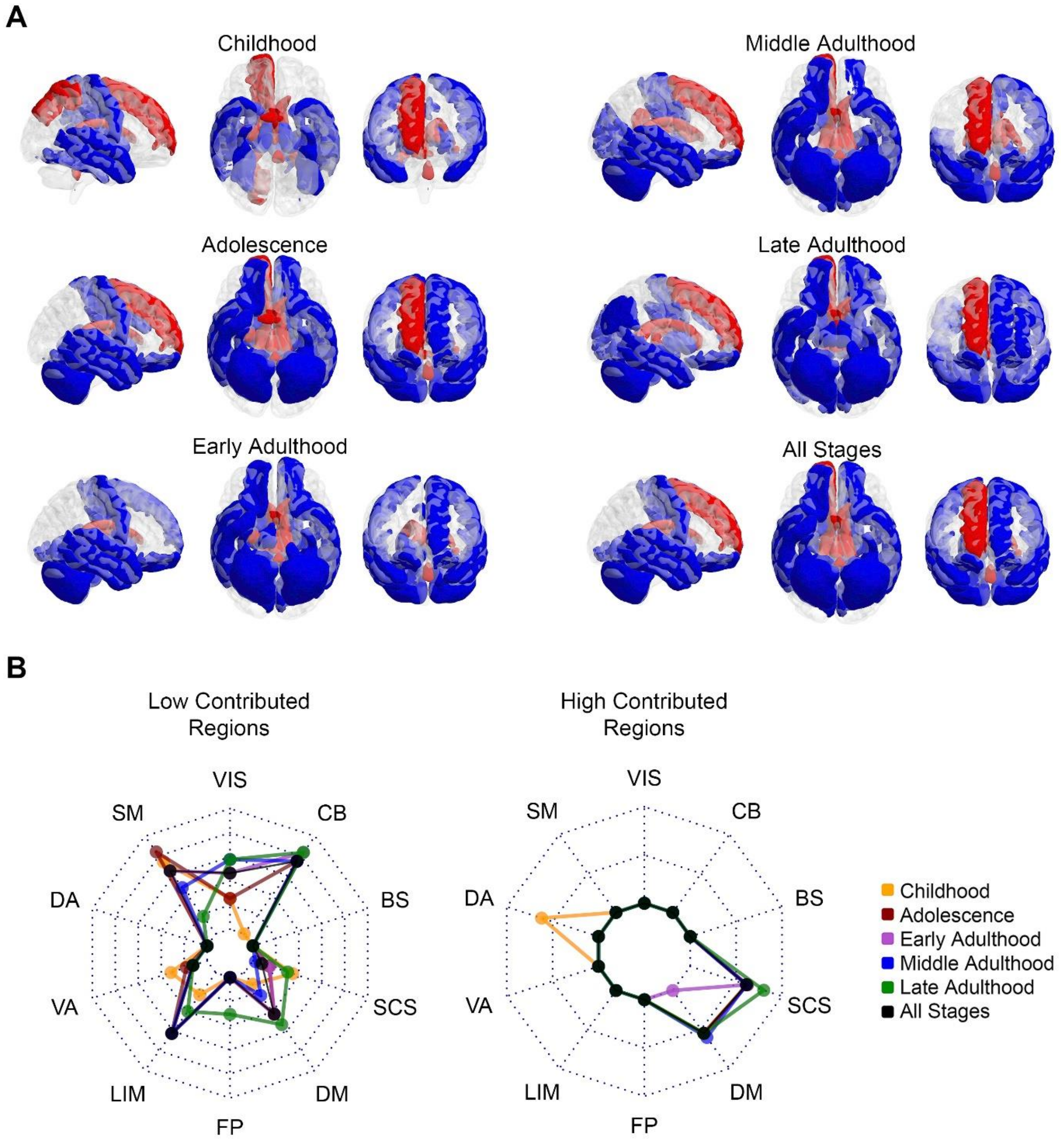


**Figure S10. The pattern of regional contribution in frustration formation for DKT parcellation.** Red-colored and blue-colored areas of the brain maps indicate DKT atlas's regions with significantly greater and significantly lower contributions in frustration formation. Brain maps demonstrate the patterns for various lifespan stages and all stages in three representational planes of sagittal, axial, and coronal. Abbreviation; VIS: visual, SM: somatomotor, DA: dorsal attention, VA: ventral attention, LIM: limbic, FP: frontoparietal, DM: default mode, SCS: subcortical structures, BS: brainstem, CB: cerebellum.

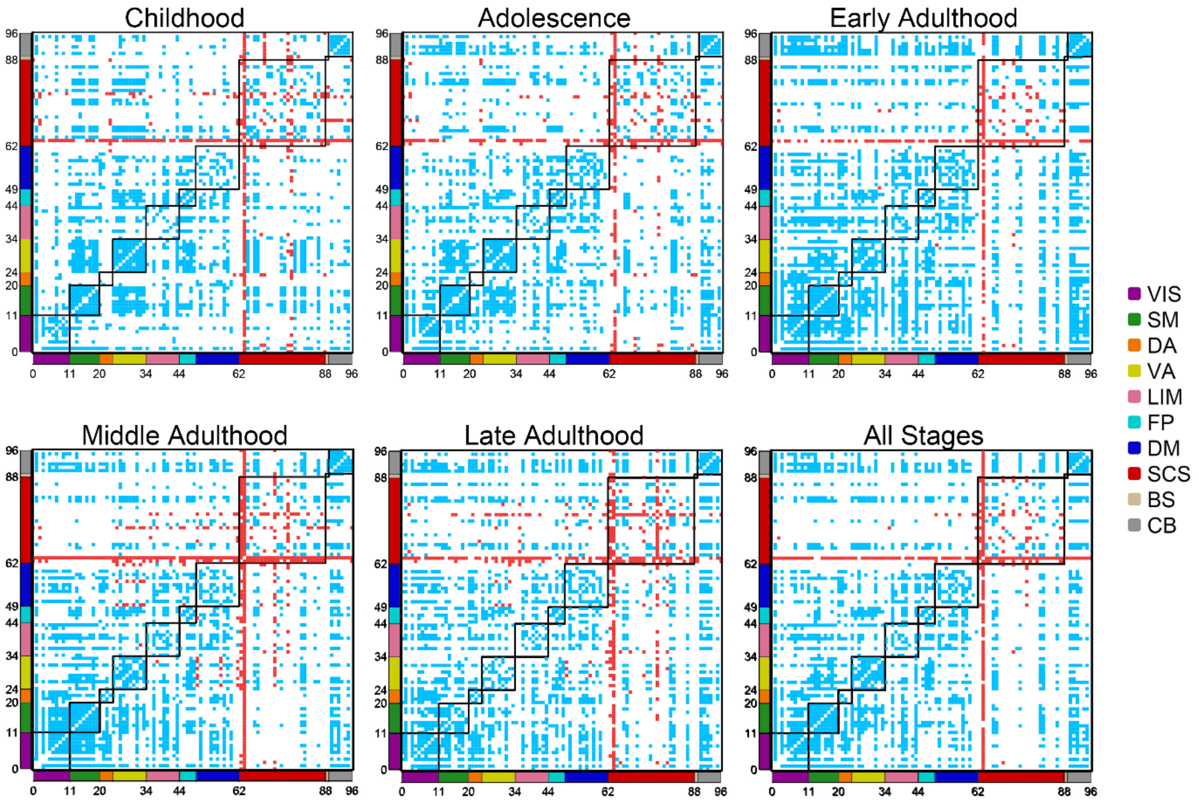


**Figure S11 - Contribution maps of functional connections in frustration formation for DKT parcellation.** The first five maps demonstrate the maps related to lifespan stages and the last one corresponds without considering the stage. Shen's 268 ROIs are categorized into 10 networks of Figure 2 by axes coloring. Cells also represent functional connections between every two ROIs. Red cells display those connections with a significantly greater contribution to frustration formation and blue cells indicate those connections with significantly lower involvement. Black squares discriminate between network connections. Abbreviation; VIS: visual, SM: somatomotor, DA: dorsal attention, VA: ventral attention, LIM: limbic, FP: frontoparietal, DM: default mode, SCS: subcortical structures, BS: brainstem, CB: cerebellum.

### Tables:

| Stage | Childhood | Adolescence | Early Adulthood | Middle Adulthood | Late Adulthood |
|---|---|---|---|---|---|
| **Visual** | 2.91E-19 (-0.846) | 1.75E-22 (-0.83) | 3.05E-38 (-0.811) | 1.63E-32 (-0.84) | 2.27E-13 (-0.869) |
| **Somatomotor** | 3.69E-20 (-0.868) | 1.68E-24 (-0.871) | 1.63E-43 (-0.866) | 9.79E-35 (-0.867) | 1.19E-12 (-0.862) |
| **Dorsal Attention** | 3.69E-20 (-0.868) | 1.68E-24 (-0.863) | 2.57E-43 (-0.866) | 1.05E-34 (-0.867) | 1.22E-13 (-0.87) |
| **Ventral Attention** | 3.69E-20 (-0.868) | 1.68E-24 (-0.871) | 4.44E-43 (-0.86) | 1.04E-34 (-0.867) | 6.47E-13 (-0.865) |
| **Limbic** | 3.70E-16 (-0.773) | 2.91E-19 (-0.765) | 1.68E-36 (-0.792) | 4.01E-34 (-0.861) | 1.39E-09 (-0.801) |
| **Frontoparietal** | 2.16E-19 (-0.849) | 4.68E-23 (-0.838) | 2.90E-34 (-0.768) | 2.15E-27 (-0.771) | 3.49E-11 (-0.84) |
| **Default Mode** | 3.69E-20 (-0.867) | 2.04E-24 (-0.863) | 5.50E-42 (-0.854) | 2.14E-34 (-0.861) | 2.27E-13 (-0.869) |
| **Subcortical Structures** | 1.22E-13 (-0.708) | 1.49E-10 (-0.555) | 7.22E-01 (0.022) | 3.55E-03 (-0.233) | 1.22E-01 (-0.271) |
| **Brainstem** | 2.74E-04 (-0.371) | 9.11E-07 (-0.434) | 1.49E-02 (-0.178) | 6.82E-08 (-0.397) | 1.17E-01 (-0.297) |
| **Cerebellum** | 2.91E-19 (-0.846) | 7.38E-22 (-0.818) | 8.84E-43 (-0.86) | 5.48E-34 (-0.854) | 4.97E-13 (-0.868) |

**Table S1. Statistics related to within-network contribution of canonical networks in frustration formation.** Corrected p-values of the Wilcoxon paired signed-rank test are presented in cells and corresponding r effect sizes are parenthesized below them. Highlighted cells denote corrected p-value lower than 0.05 with a large r effect size.

| Stage | Childhood | Adolescence | Early Adulthood | Middle Adulthood | Late Adulthood |
|---|---|---|---|---|---|
| **Visual** | 1.91E-06 (-0.482) | 1.00E-09 (-0.543) | 7.42E-22 (-0.613) | 7.06E-22 (-0.688) | 6.62E-10 (-0.82) |
| **Somatomotor** | 2.41E-19 (-0.854) | 4.01E-24 (-0.863) | 1.53E-42 (-0.86) | 7.74E-34 (-0.854) | 3.44E-08 (-0.769) |
| **Dorsal Attention** | 2.59E-19 (-0.853) | 8.34E-23 (-0.838) | 1.57E-39 (-0.823) | 1.95E-31 (-0.826) | 4.03E-08 (-0.766) |
| **Ventral Attention** | 5.07E-18 (-0.823) | 1.62E-22 (-0.83) | 3.48E-28 (-0.694) | 2.82E-19 (-0.644) | 2.85E-02 (-0.419) |
| **Limbic** | 2.20E-01 (-0.184) | 2.77E-01 (-0.15) | 4.42E-09 (-0.391) | 2.28E-18 (-0.629) | 7.86E-02 (-0.362) |
| **Frontoparietal** | 4.03E-10 (-0.613) | 1.76E-13 (-0.843) | 6.77E-13 (-0.469) | 4.27E-12 (-0.509) | 6.56E-06 (-0.673) |
| **Default Mode** | 4.98E-09 (-0.576) | 8.74E-11 (-0.573) | 1.30E-17 (-0.549) | 1.24E-11 (-0.498) | 1.99E-04 (-0.59) |
| **Subcortical Structures** | 2.85E-02 (-0.267) | 3.70E-01 (0.098) | 4.04E-13 (0.474) | 2.09E-08 (0.421) | 2.11E-01 (0.303) |
| **Brainstem** | 1.10E-07 (0.528) | 1.08E-03 (0.318) | 1.06E-07 (0.358) | 3.70E-01 (0.0917) | 2.77E-01 (0.261) |
| **Cerebellum** | 6.41E-04 (-0.365) | 3.79E-06 (-0.423) | 1.80E-26 (-0.676) | 2.47E-20 (-0.662) | 1.97E-06 (-0.699) |

**Table S2. Statistics related to between-network (type I) contribution of canonical networks in frustration formation.** Corrected p-values of the Wilcoxon paired signed-rank test are presented in cells and corresponding r effect sizes are parenthesized below them. Highlighted cells denote corrected p-value lower than 0.05 with a large r effect size.

| Stage | Childhood | Adolescence | Early Adulthood | Middle Adulthood | Late Adulthood |
|---|---|---|---|---|---|
| **Visual** | 3.15E-09 (-0.575) | 1.13E-15 (-0.69) | 1.68E-32 (-0.749) | 8.28E-32 (-0.826) | 1.89E-12 (-0.862) |
| **Somatomotor** | 1.32E-19 (-0.856) | 8.93E-24 (-0.855) | 1.72E-40 (-0.835) | 1.97E-32 (-0.84) | 2.95E-10 (-0.823) |
| **Dorsal Attention** | 4.68E-20 (-0.867) | 4.23E-24 (-0.863) | 1.38E-40 (-0.835) | 2.12E-32 (-0.84) | 2.04E-11 (-0.847) |
| **Ventral Attention** | 7.92E-19 (-0.839) | 4.23E-24 (-0.863) | 1.04E-24 (-0.651) | 1.19E-18 (-0.632) | 4.05E-05 (-0.618) |
| **Limbic** | 2.54E-12 (-0.673) | 8.01E-14 (-0.647) | 4.72E-29 (-0.706) | 7.71E-28 (-0.778) | 1.59E-05 (-0.642) |
| **Frontoparietal** | 2.45E-18 (-0.827) | 1.61E-23 (-0.847) | 9.02E-39 (-0.817) | 3.95E-34 (-0.861) | 4.26E-13 (-0.867) |
| **Default Mode** | 2.14E-15 (-0.756) | 4.89E-14 (-0.652) | 5.86E-35 (-0.774) | 1.55E-24 (-0.73) | 1.67E-08 (-0.772) |
| **Subcortical Structures** | 8.67E-10 (-0.595) | 6.38E-06 (-0.408) | 3.89E-06 (-0.311) | 3.80E-07 (-0.38) | 4.95E-03 (-0.471) |
| **Brainstem** | 1.08E-01 (0.201) | 9.35E-01 (0.083) | 9.35E-01 (0.046) | 6.20E-02 (-0.172) | 9.35E-01 (-0.013) |
| **Cerebellum** | 9.27E-14 (-0.713) | 4.85E-18 (-0.741) | 1.95E-37 (-0.805) | 7.26E-31 (-0.819) | 2.04E-11 (-0.847) |

**Table S3. Statistics related to between-network (type II) contribution of canonical networks in frustration formation.** Corrected p-values of the Wilcoxon paired signed-rank test are presented in cells and corresponding r effect sizes are parenthesized below them. Highlighted cells denote corrected p-value lower than 0.05 with a large r effect size.

| Stage | Childhood | Adolescence | Early Adulthood | Middle Adulthood | Late Adulthood |
|---|---|---|---|---|---|
| **Visual** | 1.68E-14 (0.737) | 6.12E-17 (0.723) | 9.00E-16 (0.52) | 3.96E-10 (0.464) | 1.00E+00 (0.153) |
| **Somatomotor** | 1.54E-01 (-0.229) | 3.63E-02 (-0.247) | 1.00E+00 (-0.077) | 1.00E+00 (0.01) | 4.00E-01 (0.296) |
| **Dorsal Attention** | 4.00E-01 (-0.184) | 2.03E-01 (-0.196) | 6.11E-01 (-0.109) | 1.00E+00 (-0.025) | 1.00E+00 (0.102) |
| **Ventral Attention** | 1.00E+00 (-0.021) | 1.00E+00 (0.007) | 3.27E-12 (0.456) | 1.66E-15 (0.578) | 1.54E-05 (0.663) |
| **Limbic** | 1.32E-14 (0.74) | 9.19E-21 (0.801) | 4.48E-24 (0.645) | 1.09E-16 (0.6) | 1.85E-06 (0.706) |
| **Frontoparietal** | 6.22E-05 (0.421) | 5.33E-03 (0.293) | 3.06E-13 (0.475) | 6.93E-16 (0.585) | 3.86E-01 (0.303) |
| **Default Mode** | 7.66E-12 (0.664) | 9.40E-14 (0.649) | 2.76E-27 (0.688) | 6.30E-29 (0.792) | 1.29E-06 (0.713) |
| **Subcortical Structures** | 5.18E-08 (0.545) | 2.70E-16 (0.708) | 9.76E-33 (0.749) | 2.06E-28 (0.785) | 1.24E-06 (0.714) |
| **Brainstem** | 5.54E-19 (0.846) | 1.64E-20 (0.795) | 9.57E-35 (0.774) | 5.44E-23 (0.702) | 5.55E-05 (0.634) |
| **Cerebellum** | 1.16E-12 (0.687) | 3.65E-11 (0.582) | 2.78E-03 (0.231) | 6.14E-13 (0.527) | 5.55E-05 (0.634) |

**Table S4. Statistics related to between-network (type III) contribution of canonical networks in frustration formation.** Corrected p-values of the Wilcoxon paired signed-rank test are presented in cells and corresponding r effect sizes are parenthesized below them. Highlighted cells denote corrected p-value lower than 0.05 with a large r effect size.

| | Childhood | Adolescence | Early Adulthood | Middle Adulthood | Late Adulthood |
|---|---|---|---|---|---|
| **State 1** | 1.00E+00 (0.073) | 4.79E-04 (-0.33) | 1.68E-02 (-0.187) | 7.31E-08 (-0.398) | 2.33E-01 (-0.287) |
| **State 2** | 2.72E-04 (-0.379) | 1.00E+00 (-0.047) | 1.04E-01 (0.146) | 1.00E+00 (0.046) | 5.69E-01 (-0.214) |
| **State 3** | 7.93E-04 (0.354) | 1.00E+00 (0.018) | 1.12E-01 (0.144) | 1.00E+00 (-0.052) | 7.71E-01 (0.189) |
| **State 4** | 1.00E+00 (-0.049) | 9.83E-05 (0.362) | 1.09E-02 (0.194) | 3.22E-08 (0.408) | 1.50E-01 (0.312) |

**Table S5. Statistics related to Hemisphere contribution in frustration formation.** Corrected p-values of the Wilcoxon paired signed-rank test are presented in cells and corresponding r effect sizes are parenthesized below them.

| Dataset | Study | Child (6 -12) | Adolescent (12 -18) | Early Adult (18 – 40) | Middle Adult (40 – 65) | Late Adult (65 < ) | Total Subjects |
|---|---|---|---|---|---|---|---|
| Southwest | Southwest University | - | - | 173 | 211 | 48 | 432 |
| ABIDEI | NYU Langone Medical Center | 28 | 31 | 26 | - | - | 85 |
| | San Diego State University | - | 20 | - | - | - | 20 |
| | University of Michigan | 13 | 37 | 9 | - | - | 59 |
| | University of Utah School of Medicine | 1 | 6 | 25 | - | - | 32 |
| | Yale Child Study Center | 9 | 15 | | - | - | 24 |
| ABIDEII | ETH Zürich | - | 2 | 14 | - | - | 16 |
| | Georgetown University | 36 | 12 | - | - | - | 48 |
| | NYU Langone Medical Center | 24 | 2 | 1 | - | - | 27 |
| | San Diego State University | 10 | 12 | - | - | - | 22 |
| | Trinity Centre for Health Sciences | - | 10 | 5 | - | - | 15 |
| | University of Utah School of Medicine | - | 1 | 12 | - | - | 13 |
| **Total Subjects (Female)** | | 121 (32) | 148 (32) | 265 (117) | 211 (140) | 48 (29) | 793 (350) |
| **Age (Mean ± SD)** | | 9.7 ± 1.4 | 14.8 ± 1.69 | 25.04 ± 4.8 | 54.06 ± 6.53 | 71.31 ± 3.83 | 31.31 ± 19.78 |

**Table S6. Subject demographics.**

| Dataset | Study | Final Selected Subjects | Voxel Size(mm) | Structural Flip Angle (Deg) | Functional Flip Angle (Deg) | Structural Echo Time (ms) | Functional Echo Time (ms) | Structural Repetition Time (ms) | Functional Repetition Time (ms) |
|---|---|---|---|---|---|---|---|---|---|
| Southwest | Southwest University | 159 | 1.0×1.0×1.0 | 9 | 90 | 2.52 | 30 | 1900 | 2000 |
| ABIDEI | NYU Langone Medical Center | 61 | 1.3×1.0×1.3 | 7 | 90 | 3.25 | 15 | 2530 | 2000 |
| | San Diego State University | 15 | - | 8 | 90 | min full | 30 | 600 | 2000 |
| | University of Michigan | 46 | 3.438×3.438×3.0 | 15 | 90 | 1.8 | 30 | 500 | 2000 |
| | University of Utah School of Medicine | 32 | 1.0×1.0×1.2 | 9 | 90 | 2.91 | 28 | 2300 | 2000 |
| | Yale Child Study Center | 17 | 1.0×1.0×1.0 | 9 | 60 | 1.73 | 25 | 1230 | 2000 |
| ABIDEII | ETH Zürich | 16 | 0.898×0.898×0.899 | 8 | 90 | shortest | 25 | 8.4 | 2000 |
| | Georgetown University | 25 | 1.0×1.0×1.0 | 7 | 90 | 3.5 | 30 | 2530 | 2000 |
| | NYU Langone Medical Center | 26 | 1.3×1.0×1.3 | 7 | 90 | 3.25 | 15 | 2530 | 2000 |
| | San Diego State University | 20 | - | 8 | 90 | min full | 30 | 600 | 2000 |
| | Trinity Centre for Health Sciences | 15 | 0.898×0.898×0.899 | 8 | 90 | shortest | 27 | 8.4 | 2000 |
| | University of Utah School of Medicine | 11 | 1.0×1.0×1.2 | 9 | 90 | 2.91 | 28 | 2300 | 2000 |

**Table S7. Study-specific scan parameters.**
